# Preimplantation factor (PIF) links embryo-derived signaling to maternal pancreatic β-cell adaptation through an ERα-dependent pathway

**DOI:** 10.64898/2026.09.10.750557

**Authors:** R. Pascua-Maestro, A. Sartorius-Yáñez, S. Hernández de la Red, T. Boronat-Belda, M. Arévalo-Martínez, M. Martín-Martín, M. Royo, M. López-Corrales, N. Pascual, M.-P. Marco, M. Daniel-Mozo, C Broca, G Perdomo, I Cózar-Castellano, P. Alonso-Magdalena, B. Merino

## Abstract

Pregnancy requires maternal pancreatic β-cells adaptations to increased insulin demand, yet the embryo-derived signals contributing to this response remain poorly defined. Here we identify preimplantation factor (PIF), an embryo-derived peptide present in maternal circulation from early gestation, as a regulator of β-cell adaptation. In a murine model of gestational diabetes mellitus (GDM), circulating PIF levels were reduced during mid-gestation, indicating that this endogenous signal is altered under gestational metabolic dysfunction. Conversely, chronic exposure of non-pregnant female mice to synthetic PIF (sPIF) recapitulated key temporal features of gestational β-cell adaptation, including early β-cell proliferation, increased glucose-stimulated circulating C-peptide, expansion of β-cell mass and sustained enhancement of *ex vivo* glucose-stimulated insulin secretion (GSIS). Mechanistically, sPIF activated rapid ERK-, AKT- and PKA-dependent signaling that converged on estrogen receptor alpha (ERα) phosphorylation and nuclear translocation. Pharmacological inhibition and genetic silencing demonstrated that ERα is required for full propagation of the functional and kinase responses. This kinase–ERα axis was conserved in human islets, where sPIF enhanced insulin secretion in an ERα-dependent manner. These findings identify PIF as an embryo-derived metabolic signal supporting maternal β-cell compensation and suggest that reduced PIF availability may contribute to inadequate β-cell adaptation in GDM.

## 1. Introduction

Preimplantation factor (PIF) is a 15-amino acid peptide (MVRIKPGSANKPSDD) secreted by the mammalian embryo and placenta, detectable in maternal circulation from the earliest stages of gestation and maintained throughout viable pregnancy [1–5]. PIF has been primarily characterized as a regulator of embryo–maternal dialogue, with reported effects on immune tolerance, trophoblast invasion, cell adhesion, tissue remodeling and apoptosis [4–7]. The synthetic analogue of PIF (sPIF) reproduces the biological activity of the native peptide *in vitro* and *in vivo* and has shown a favorable safety profile in clinical settings [8].

Pregnancy represents a physiological state of increased metabolic demand and progressive insulin resistance. To maintain maternal glucose homeostasis and ensure an adequate nutrient supply to the fetus, pancreatic β-cells undergo a coordinated adaptive response that includes increased insulin synthesis, enhanced glucose-stimulated insulin secretion (GSIS) and expansion of β-cell mass [9–11]. Failure of this compensatory response contributes to metabolic dysregulation during pregnancy and is associated with gestational diabetes mellitus (GDM) [12–13]. Although several hormonal and nutrient-derived signals have been implicated in this process, including estrogens, progesterone, prolactin, placental lactogens, cortisol and metabolic substrates, the contribution of embryo-derived circulating factors to maternal β-cell adaptation remains unexplored. Given its early and sustained presence in maternal circulation, PIF represents a candidate signal linking embryonic development to maternal β-cell physiology. Consistent with this possibility, our previous work demonstrated that chronic sPIF treatment enhances β-cell function and improves glucose tolerance in a preclinical model of diet-induced obesity, suggesting that PIF may regulate pancreatic endocrine function under conditions of increased metabolic demand [14]. These findings raised the possibility that the effects of sPIF on insulin secretion reflect a broader role in maternal–fetal metabolic communication.

Here, we investigated the role of PIF in female glucose metabolism and β-cell adaptation. We determined whether circulating PIF levels are altered in a validated murine model of GDM and whether sustained sPIF exposure promotes functional and structural β-cell adaptations resembling those observed during pregnancy. We further investigated the signaling mechanisms underlying this response, focusing on estrogen receptor-associated pathways and their functional relevance in murine and human islets.

## 2. Methods

### 2.1 Human islets

Experiments involving human islets obtained from Montpellier Hospital (LTCD) were conducted in accordance with the local ethics committee (Biological Resources Center, Collection IRB 5 “Human Islets of Langerhans”, Biobank no. BB-0033-00031, CHU Montpellier), the institutional ethics committee of the French Agence de la Biomédecine (ABM no. PFS 13-008), and the French Ministry of Research (DC-2011-1401 and AC-2017-3039). Human pancreata were obtained from brain-dead non-diabetic donors following appropriate ethical approval, consent procedures and tissue handling regulations. The human islets used in this study were obtained from two non-diabetic female donors, aged 43 and 70 years, with glycated hemoglobin values within the normal range.

Isolated islets were obtained according to a slightly modified version of the automated method of C. Ricordi, as previous described [14]. Briefly, following collagenase digestion at 37 °C in a digestion chamber, human islets were purified in a continuous density gradient using a COBE 2991 cell processor. Following isolation, human islets were cultured for recovery for 3 days at 37 °C, in a 5% CO₂ atmosphere, in CMRL 1066 medium (Life Technologies,USA) containing 5.6 mM glucose supplemented with 7.5% FBS, 2 mM glutamine, 25 mM HEPES, and antibiotics. The islet purity was 85–90%, the viability was 80%, and the islet size index was 1.

### 2.2 Mice models

12 weeks female C57BL/6J mice were purchased form Envigo (C57BL/6JOlaHsd; Barcelona, Spain). The mice were housed in ventilated cages under a 12-hour:12-hour light–dark cycle with water available ad libitum, at the animal facility of the University of Valladolid (UVa). The Animal Care and Use Committee of the UVa approved all experiments (protocol #13504003). Randomization of mice was performed before the treatment with saline or sPIF using an Excel spreadsheet. Then mini-pumps (Alzet, USA) containing saline or sPIF were subcutaneously implanted in the upper back of the mice. The rate of delivery was 1 mg/kg/day. The pumps remained implanted for 21 days (time-pregnancy window). Afterwards, mice were euthanized, and blood and tissues (pancreas and pancreatic islets) were collected for plasma biochemistry analyses, immunohistochemistry and GSIS assessment.

For pregnant mice models female C57BL/6J mice (C57BL/6JOlaHsd; 9–10 weeks of age) were obtained from Envigo (Barcelona, Spain). Animals were housed under controlled environmental conditions with ad libitum access to food and water throughout the study. All experimental procedures regarding this model were approved by the Animal Ethics Committee of Miguel Hernandez University and the competent regional authority (Generalitat Valenciana; approval IDs: 2019/VSC/PEA/0243 and 2024-VSC-PEA-0194). All experiments were conducted in accordance with Directive 2010/63/EU of the European Parliament on the protection of animals used for scientific purposes.

The GDM animal model was established as previously described [15]. Briefly, female mice were randomly assigned to receive either a standard control diet (10% of total energy derived from fat; D12450J, Research Diets, USA) or a high-fat diet (HFD; 60% of total energy derived from fat; D12492, Research Diets, USA). Dietary intervention was initiated three days before mating and maintained throughout pregnancy. Females were housed overnight with males, and successful mating was confirmed by the presence of a vaginal plug the following morning, which was designated as gestational day (GD) 0. Blood samples were collected from the saphenous vein in the non-fasted state one day before the initiation of the dietary intervention (baseline) and on GD8, GD15, and GD18. On GD18, pregnant mice were euthanized by decapitation in the non-fasted state.

### 2.3 Mouse islet isolation and *ex vivo* GSIS

Mouse islets isolation, assessment of GSIS and quantification of intracellular insulin release were performed as previously described [14].

### 2.4 Min-6 β-cells culture, isolated islets culture and treatments

Min 6 β-cells were grown in Dulbecco’s modified Eagle medium (4.5 g/L glucose, Gibco, USA) supplemented with 15% fetal bovine serum (Gibco, Brazil), 100 U/mL penicillin (Gibco, USA), 100 μg/mL streptomycin (Gibco, USA), 1 mM sodium pyruvate (Gibco, USA) and 50 μM β-mercaptoethanol (Sigma,USA) at 37 °C and 5% CO₂ in a humidified atmosphere to 80% confluence. Twenty-four hours before treatments and functional assays, the culture medium was replaced with phenol red–free medium of identical composition supplemented with charcoal-stripped fetal bovine serum (Gibco, Brazil). These conditions were used to minimize the presence of estrogenic compounds in the culture medium and to avoid potential confounding effects on estrogen-dependent signaling pathways.

For isolated islets, culture and treatments were carried out in phenol red-free RPMI 1640 (Gibco, USA) supplemented with 10% charcoal-stripped fetal bovine serum (Gibco, USA), 5.5 mM glucose (Sigma-Aldrich, USA), 100 U/mL penicillin and 100 μg/mL streptomycin (Gibco, USA). Cells and islets were treated with synthetic preimplantation factor (sPIF, 50 nM; synthesized in-house, see below), 17β-estradiol (E2, 10 nM; Sigma-Aldrich, USA), the selective ERα antagonist MPP (10 nM; Tocris Bioscience, United Kingdom) and the estrogen receptor downregulator ICI 182,780 (1 µM; Tocris Bioscience, United Kingdom). E2 and the specific inhibitors were used at experimentally selected concentrations informed by previous studies investigating estrogen receptor signaling and β-cell biology [16-19.]. All compounds were dissolved in the appropriate vehicle, which was included in control conditions at equivalent concentrations.

### 2.5 siRNA-mediated ERα knockdown

MIN6 β-cells were transfected with an ON-TARGETplus SMARTpool siRNA targeting mouse **ERα/Esr1** (Dharmacon, USA) using Lipofectamine 2000 Transfection Reagent (Invitrogen, USA), according to the manufacturer’s instructions. Cells were exposed to the transfection mix for 6 h and then cultured in complete medium for 48 h before experimentation. An ON-TARGETplus non-targeting siRNA pool (Dharmacon, USA) was used as control. After 48 h, ERα knockdown efficiency was assessed by Western blot, and cells were treated with sPIF as indicated for downstream signaling experiments.

### 2.6 *In vitro* GSIS in MIN-6 β-cells, isolated mouse and human islets

For GSIS assays, MIN6 β-cells were preincubated in Hanks’ balanced salt solution (HBSS) supplemented with 3 mM glucose and 0.1% bovine serum albumin (BSA, Sigma, USA) for 1 h at 37 °C to stabilize basal insulin secretion. Cells were then incubated for 1 h in HBSS containing either low glucose (3 mM) or stimulatory glucose (16 mM) in the presence or absence of the indicated treatments.

For isolated islets, GSIS was performed using Krebs–Ringer bicarbonate (KRB) buffer supplemented with 0.1% BSA (Sigma, USA). Islets were preincubated for 1 h at 37 °C in KRB containing 3 mM glucose and subsequently incubated for 1 h in KRB under low (3 mM) or high (16 mM) glucose conditions, with or without treatments, as indicated. At the end of the incubation period, supernatants were collected and centrifuged to remove debris. Insulin secretion was determined using a commercially available mouse insulin ELISA kit (Crystal Chem, USA) according to the manufacturer’s instructions. Data was normalized to islet number or total DNA and expressed as fold change relative to basal conditions.

GSIS was also expressed as “Fold-change”, which was calculated as: (high-glucose insulin secretion - low-glucose insulin secretion) / low-glucose insulin secretion.

### 2.7 Intraperitoneal glucose tolerance (IpGTT) test and C-peptide

To evaluate *vivo* glucose homeostasis in our non-pregnant model treated with sPIF, we performed an intraperitoneal glucose tolerance test (IpGTT). 15 days after mini-pump implantation, mice were fasted for 16 hours and then injected with 2g glucose/kg body weight, as previously described [14]. Blood samples were collected from the tail vein using capillary tubes precoated with potassium-EDTA (Sarstedt, Germany), and plasma was obtained by centrifugation at 3300 ×g for 15 minutes at 4°C. C-peptide levels were determined by ELISA kit (Crystal-Chem, USA), following the manufacturer’s instructions.

IpGTT in pregnant mice were performed as previously described performed on GD15 to assess glucose tolerance [15]. Mice were fasted for 6 h (08:00–14:00) with free access to water. Glucose was then administered by intraperitoneal injection at a dose of 1.5 g/kg body weight. Blood was sampled from the tail vein for glucose measurements during the IpGTT. Blood glucose concentrations were measured using an automatic glucometer (Accu-Chek Compact Plus, Roche, Madrid, Spain) immediately before glucose administration (0 min) and at 10, 20, 30, 60, and 120 min after glucose injection.

### 2.8 Pancreas histomorphometry, β-cell mass and proliferation

Paraffin-embedded pancreatic sections (5 μm) were processed for immunofluorescence analysis of β-cell proliferation as previously described [20]. Briefly, sections were deparaffinized, rehydrated, subjected to antigen retrieval in citrate buffer (pH 6.0), blocked with 5% normal donkey serum and incubated overnight at 4 °C with primary antibodies against insulin (1:1000; Invitrogen, USA) and Ki67 (1:500; Abcam, United Kingdom). Sections were then incubated with species-appropriate Alexa Fluor-conjugated secondary antibodies, including Alexa Fluor 488 for Ki67 and Alexa Fluor 594 for insulin, and mounted with Fluoroshield™ containing DAPI (Sigma-Aldrich, USA). β-cell proliferation was quantified as the percentage of insulin⁺/Ki67⁺ cells over total insulin⁺ cells, with more than 1000 insulin-positive cells evaluated per section. Pancreatic histomorphometry, including β-cell area, β-cell mass, islet number and islet size, was performed in a double-blind manner as previously described [20]. For each pancreas, three sections separated by 200 μm were stained and analyzed. β-cell mass was calculated by measuring the insulin-positive area relative to total pancreatic area using image analysis software and multiplying by pancreatic weight.

### 2.9 Immunofluorescence and confocal microscopy

Paraffin-embedded pancreatic sections were deparaffinized in xylene, rehydrated through a graded ethanol series and washed in PBS. For endocrine hormone immunofluorescence analysis, antigen retrieval was performed in citrate buffer (pH 6.0) for 30 min using a pressure cooker. After washing PBS, sections were blocked for 1 h at room temperature in PBS containing 10% fatty acid-free BSA, 1% normal goat serum and 0.05% Triton X-100. Sections were then incubated overnight at 4 °C with primary antibodies against insulin (mouse monoclonal anti-insulin, Sigma-Aldrich, I2018; 1:1000), glucagon (rabbit anti-glucagon, Abcam, ab92517; 1:2000) and somatostatin (rat anti-somatostatin, Abcam, ab30788; 1:250) diluted in blocking solution. After PBS washes, sections were incubated with species-appropriate Alexa Fluor-conjugated secondary antibodies (Invitrogen), including Alexa Fluor 488, Alexa Fluor 594 and Alexa Fluor 647, diluted in blocking solution. Sections were washed in PBS and mounted with Fluoroshield™ containing DAPI (Sigma-Aldrich, F6057). Confocal images were acquired using a 60× oil immersion objective on a Leica DMI 6000B microscope equipped with a TCS SP5 confocal system. Fluorophores were excited using a white light laser and a 405 nm laser line, controlled by LAS AF software. Emission signals were collected using the AOBS system and spectral detectors. Images were processed and analyzed using FIJI software.

Nuclear colocalization was quantified using Pearson’s correlation coefficient and Manders’ M1 coefficient in FIJI. Nuclear regions of interest were defined from the DAPI channel and used as masks. Pearson’s correlation coefficient was calculated as the covariance between the pixel intensities of the ERα or phospho-ERα channel and the nuclear mask channel, divided by the product of their standard deviations. Thus, Pearson’s coefficient estimates the degree of spatial correlation between both signals, with values ranging from −1 to +1, where +1 indicates complete positive correlation, 0 indicates no correlation and −1 indicates inverse correlation. Manders’ M1 coefficient was calculated as the fraction of total ERα or phospho-ERα fluorescence intensity overlapping with the nuclear mask. In this analysis, M1 represents the proportion of ERα or phospho-ERα signal localized within the nuclear compartment, independently of the relative intensity distribution of the nuclear signal. Both coefficients were calculated from randomly acquired fields and used for statistical analysis as indicated in the figure legends.

### 2.10 sPIF synthesis and anti-PIF polyclonal antibody

sPIF peptide was synthesized manually using a Fmoc/tBu solid-phase strategy on a 2-chloro trityl chloride resin (CTC) at the Synthesis of Peptides Unit (U3; https://www.nanbiosis.es/portfolio/u3-synthesis-of-peptides-unit/) of the ICTS NANBIOSIS at the IQAC, as previous described [14].

To produce antibodies against PIF, the synthetic peptide was conjugated to the HCH carrier protein using the heterobifunctional crosslinker succinimidyl 3-(bromoacetamido)propionate (SBAP). Briefly, the N-hydroxysuccinimide (NHS) ester group of SBAP reacted with the primary amino groups of the peptide under mildly alkaline conditions (pH 7.5), generating a peptide-SBAP intermediate bearing a reactive bromoacetyl group. This intermediate was subsequently coupled to the carrier protein through sulfhydryl groups, yielding a stable thioether linkage and the final PIF-SBAP-HCH immunogen.

Polyclonal antibodies were produced according to the immunization protocol described by Ballesteros et al. [21]. New Zealand White rabbits (1–2 kg) were immunized intradermally on day 0 with 100 μg of PIF-SBAP-HCH conjugate emulsified (1:1, v/v) in Freund’s Complete Adjuvant. Booster immunizations containing the same amount of immunogen emulsified in Freund’s Incomplete Adjuvant were administered at monthly intervals for six months. Blood samples were collected from the marginal ear vein 10 days after each booster injection to monitor the immune response. At the end of the immunization schedule, animals were terminally bled, and blood was collected into serum-separation tubes. After clot formation, sera were recovered by centrifugation, supplemented with 0.02% (w/v) sodium azide, aliquoted and stored at −80 °C until use. Antibody production was monitored by indirect ELISA using serial dilutions of the antisera on microplates coated with the homologous conjugate.

The production of the polyclonal antisera was carried out at the Custom Antibody Service (CAbS), Unit U2 of the ICTS NANBIOSIS (IQAC-CSIC, Barcelona, Spain). All animal procedures were performed in accordance with the applicable institutional guidelines and European and national regulations governing the use of animals for scientific purposes. The experimental validation for sPIF and endogenous PIF detection is included in supplementary figure 1.

### 2.11 Western blot analysis

Protein expression and phosphorylation levels were analyzed by Western blot as previously described [22]. MIN6 β-cells and isolated human islets were lysed in ice-cold protein lysis buffer supplemented with protease and phosphatase inhibitors (Sigma-Aldrich, USA). Protein concentration was determined using a colorimetric assay, and equal amounts of protein were separated by SDS-PAGE and transferred onto PVDF membranes (Bio-Rad, USA). Membranes were blocked and incubated overnight at 4 °C with primary antibodies against the indicated proteins, as detailed in Supplementary Table 1. Phosphorylated and corresponding total proteins were detected either sequentially on the same membrane after striping or on parallel membranes loaded with equal amounts of the same protein samples. When parallel membranes were used, each membrane was analyzed with its corresponding loading control. After washing, membranes were incubated with the appropriate HRP-conjugated secondary antibodies and developed using an enhanced chemiluminescence detection system. Protein bands were visualized using a digital imaging system and quantified by densitometric analysis using FIJI software (National Institutes of Health, USA). Phosphorylated and total protein signals were independently normalized to the loading control obtained from the corresponding membrane. Phosphorylation indices were subsequently calculated as the ratio between the normalized phosphorylated and total protein signals.

### 2.12 Dot blot analysis

Dot blot assays were performed on nitrocellulose membranes to assess circulating PIF levels in plasma samples from control pregnant mice and mice from the gestational diabetes-like model. In brief, membranes were pre-equilibrated in TBS, mounted in the dot blot system and rehydrated with PBS before sample loading. Protein samples were directly applied onto the membrane (1-5 μg total protein per well), followed by two washes with PBS. Membranes were then processed as for immunoblotting, including blocking in milk and overnight incubation at 4 °C with a rabbit anti-sPIF primary antibody. After washing, membranes were incubated with the corresponding HRP-conjugated secondary antibody, and signal was detected by chemiluminescence.

### 2.14 Statistical analysis

Statistical analysis was performed using Prism v. 6.0 (GraphPad Software, Inc., San Diego, CA). The normality of the distribution of data was checked with the Kolmogorov-Smirnov test. For parametric data, comparisons between the two groups were assessed using the paired or unpaired Student’s t-test. Comparisons between more than two groups were performed using one-way or two-way analysis of variance (ANOVA) followed by Bonferroni’s post hoc test. For non-parametric data, comparisons between the two groups were assessed using the Mann-Whitney U test. Data are presented as the means ± SEM. Differences were considered significant at p less than 0.05.

## 3. Results

### 3.1 Circulating PIF is reduced in GDM and chronic sPIF enhances β-cell secretory responsiveness in females

To determine whether circulating PIF is modulated during pregnancy under metabolic stress, we first used a murine model of GDM and collected plasma samples at different stages of gestation (Fig. 1A). As expected, GDM pregnant mice showed impaired glucose tolerance compared with control pregnant mice (Fig. 1B-C). Interestingly, circulating PIF levels were altered during gestation. Plasma PIF concentrations were reduced by GDM group at gestational days 8 and 15, with no differences on days 0 and 18 (Fig. 1D-E). Consistently, the AUC of circulating PIF across gestation was significantly lower in GDM mice (Fig. 1F). These data indicate that the endogenous PIF profile is modified during pregnancy in the context of impaired maternal glucose tolerance.

**Figure 1.**
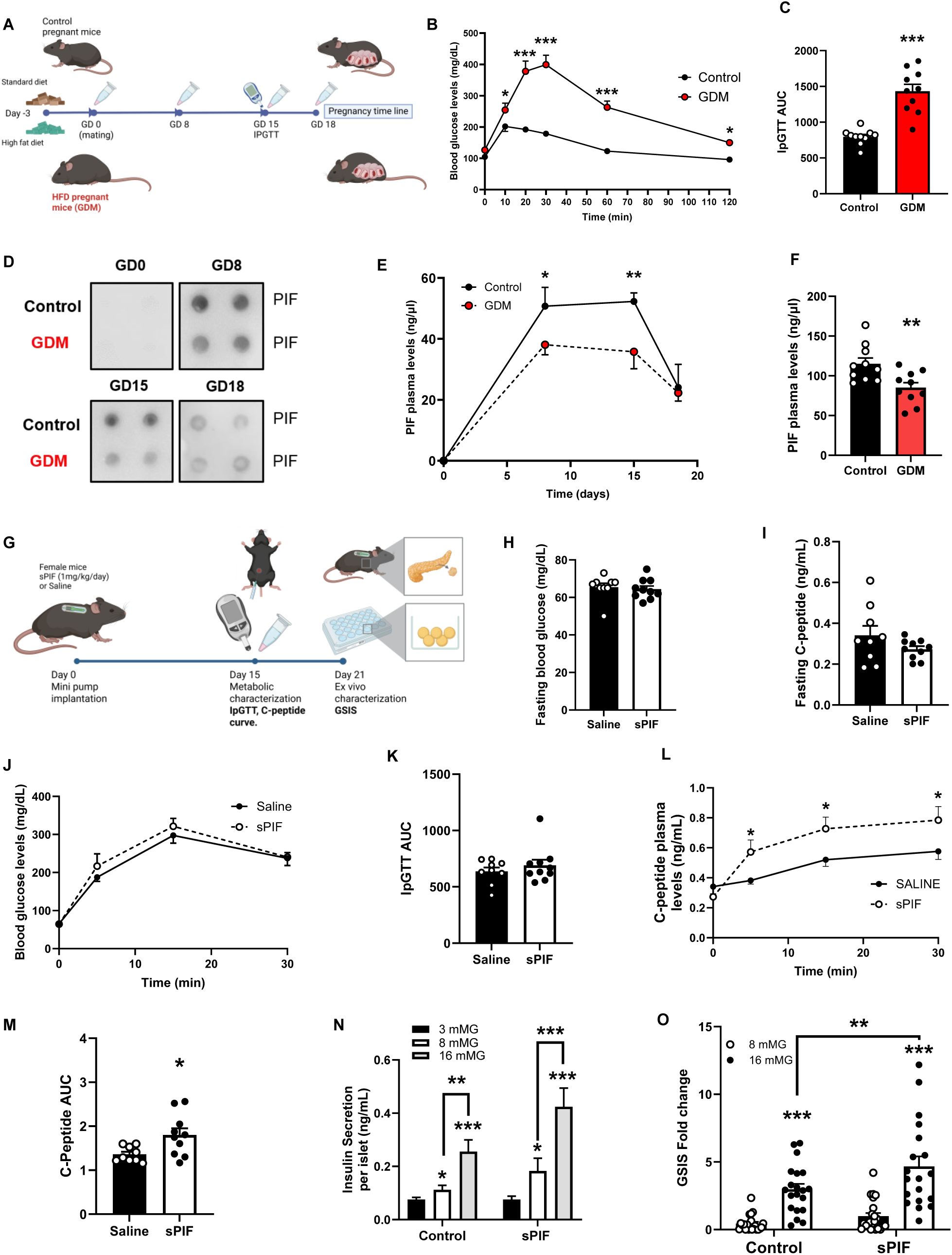
Circulating PIF is altered in GDM and chronic sPIF enhances β-cell secretory responsiveness in females. **(A)** Schematic representation of the murine pregnancy model used to compare control pregnant mice and pregnant mice exposed to a gestational diabetes-like metabolic challenge. Plasma samples were collected at gestational days 0, 8, 15 and 18.**(B-C)** IpGTT from control pregnant mice vs. GDM and corresponding area under curve (AUC). **(D)** Representative dot blot of plasma PIF levels. Each point represent one single sample in one specific time point. **(E-F)** Plasma PIF levels measured at the indicated gestational time points and AUC (10 animals per group). **(G)** Schematic representation of the experimental model of continuous sPIF administration in non-pregnant female mice using subcutaneous osmotic pumps. *In vivo* metabolic characterization was performed after 15 days of treatment, and *ex vivo* GSIS was assessed in isolated islets after 21 days of treatment.**(H)** Fasting blood glucose levels in saline- and sPIF-treated mice.**(I)** Fasting plasma C-peptide levels.**(J)** Blood glucose levels during IpGTT.**(K)** AUC of the glucose curve during the IPGTT.**(L)** Plasma C-peptide levels during the IpGTT. **(M)** AUC of the C-peptide curve during the IpGTT. 9-10 animals per group **(N)** *Ex vivo* GSIS in islets isolated from saline- and sPIF-treated mice after 21 days of treatment. **(O)** Fold change in GSIS calculated from the *ex vivo* GSIS. 20-21 islet preparations from 4 animals per condition. Data are presented as mean ± SEM. Each dot represents one animal or one independent islet preparation, as indicated. Statistical analysis was performed using unpaired Student’s t-test or two-way ANOVA followed by Bonferroni’s post hoc test, as appropriate. * p < 0.05, **p < 0.01, *** p<0.001.

We next evaluated whether sustained exposure to sPIF could modulate β-cell function in non-pregnant female mice. To this end, females were treated continuously with sPIF using subcutaneous osmotic pumps. *In vivo* metabolic characterization was performed after 15 days, and islets were isolated after 21 days to assess *ex vivo* GSIS (Fig. 1G). Fasting blood glucose levels were not different between saline- and sPIF-treated mice (Fig. 1H). Similarly, fasting plasma C-peptide levels remained unchanged (Fig. 1I). In this line, glucose tolerance evaluated by ipGTT and the corresponding AUC were not significantly modified by sPIF treatment (Fig. 1J-K).

Despite the absence of changes in glucose tolerance, sPIF-treated mice displayed a clear enhancement of the secretory response to glucose *in vivo*. Plasma C-peptide levels during the ipGTT were increased in sPIF-treated females, particularly after glucose overload (Fig. 1L-M). These results indicate that chronic sPIF exposure enhances β-cell secretory responsiveness during a glucose challenge without altering fasting glycaemia.

To determine whether this enhanced secretory response reflected intrinsic changes in islet function, GSIS was evaluated *ex vivo* in islets isolated from mice treated for 21 days. Islets from sPIF-treated females showed increased insulin secretion in response to glucose stimulation compared with saline-treated controls (Fig. 1N). This effect was also reflected in a higher fold change in GSIS (Fig. 1O). Together, these findings indicate that chronic sPIF exposure enhances β-cell responsiveness to glucose both *in vivo* and *ex vivo*, supporting the idea that sPIF induces a sustained functional adaptation of the islet.

### 3.2 sPIF induces β-cell expansion and early proliferative responses

To determine whether the enhanced β-cell secretory response induced by sPIF was associated with structural changes in the endocrine pancreas, pancreatic sections from saline- and sPIF-treated female mice were analyzed after 21 days of treatment (Fig. 2A). Morphometric analysis revealed a significant increase in relative β-cell area in sPIF-treated mice compared with saline-treated controls (Fig. 2B). Consistently, β-cell mass was also increased after chronic sPIF treatment (Fig. 2C), indicating that sustained exposure to PIF peptide promotes expansion of the β-cell compartment.

**Figure 2.**
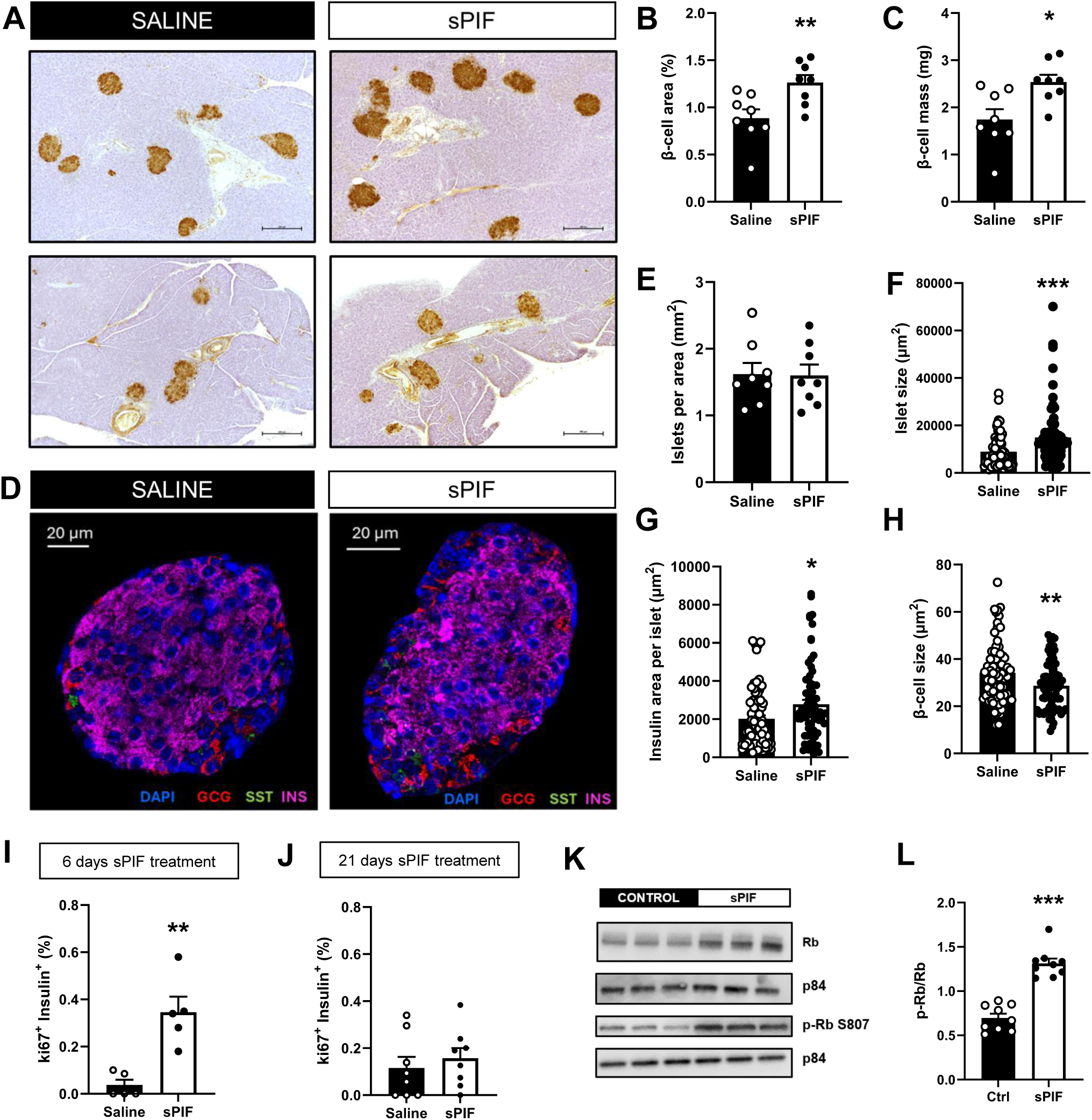
sPIF increases β-cell mass and islet size by inducing an early proliferative response. **(A)** Representative pancreatic sections from saline- and sPIF-treated female mice after 21 days. **(B)** Quantification of relative β-cell area, expressed as insulin-positive area relative to total pancreatic area.**(C)** β-cell mass calculated from insulin-positive area and pancreatic weight. 8 animals per condition. **(D)** Representative immunofluorescence images of pancreatic islets stained for insulin, glucagon and somatostatin.**(E)** Quantification of islet number normalized to pancreatic area.**(F)** Mean islet size in saline- and sPIF-treated mice.**(G)** Insulin-positive area per islet.**(H)** β-cell size calculated from insulin-positive area and β-cell number. 75-77 islets from 8 animals per condition **(I)** Quantification of Ki67-positive β-cells after 6 days of sPIF treatment, expressed as the percentage of insulin-positive cells co-expressing Ki67. 5 animals per condition. **(J)** Quantification of Ki67-positive β-cells after 21 days of sPIF treatment. 8 animals per condition.**(K)** Representative Western blots showing phosphorylated Rb and total Rb levels and P84 as loading control in MIN6 β-cells after 24 h of sPIF treatment.**(L)** Quantification of phosphorylated Rb, total Rb and the phospho-Rb/Rb ratio shown in panel K. 3 independent experiments in triplicate. Data are presented as mean ± SEM. Each dot represents one animal, islet or cell. Statistical analysis was performed using unpaired Student’s t-test after assessment of normality. *p< 0.05, **p< 0.01, ***p<0.001.

To further characterize islet remodeling, pancreatic sections were analyzed by immunofluorescence staining for insulin, glucagon and somatostatin (Fig. 2D). The number of islets normalized to pancreatic area was not significantly modified by sPIF treatment (Fig. 2E). However, mean islet size was increased in sPIF-treated mice (Fig. 2F), together with a higher insulin-positive area per islet (Fig. 2G). In contrast, β-cell size was reduced (Fig. 2H), suggesting that the increase in β-cell area and mass was not driven by β-cell hypertrophy, but rather by an increase in β-cell number.

We next assessed whether β-cell proliferation contributed to this phenotype. Immunofluorescence analysis of insulin and Ki67 showed that sPIF increased the percentage of Ki67-positive β-cells after 6 days of treatment (Fig. 2I). This proliferative response was no longer detected after 21 days of treatment (Fig. 2J), indicating that sPIF induces an early and transient proliferative phase that precedes the later expansion of β-cell mass.

Given the early increase in β-cell proliferation, we analyzed the activation of Rb, a key cell-cycle regulator. Western blot analysis in sPIF-treated MIN6 β-cells showed that sPIF increased phosphorylated Rb and total Rb levels after 24 h of treatment (Fig. 2K). Quantification confirmed an increase in phospho-Rb, total Rb and the phospho-Rb/Rb ratio (Fig. 2L), supporting activation of the Rb-dependent proliferative arm of the response.

Together, these findings indicate that chronic sPIF treatment promotes structural β-cell adaptation characterized by increased β-cell area, β-cell mass and islet size. The early and transient increase in Ki67-positive β-cells, together with activation of Rb signaling, supports a model in which sPIF-induced β-cell expansion is linked to proliferative mechanisms rather than β-cell hypertrophy.

### 3.3 sPIF enhances GSIS through ERα-dependent signaling in mouse islets and MIN6 β-cells

Because estrogen receptor signaling contributes to β-cell adaptation and insulin secretion during states of increased metabolic demand, including pregnancy, we next assessed whether sPIF functionally converges with estrogenic pathways [23]. To evaluate the involvement of estrogen receptor signaling in the effects of sPIF on β-cell function, GSIS was first assessed in isolated mouse islets after 24h of treatment. Both sPIF and E2 increased GSIS compared with control conditions, indicating that sPIF can enhance GSIS in a comparable manner to classical estrogenic stimulus (Fig. 3A). However, the combined treatment with sPIF and E2 did not further potentiate the response and instead showed a partial attenuation of the secretory effect, consistent with convergence onto shared signaling pathways (Fig. 3A). This pattern was also reflected in the corresponding fold change (Fig. 3B). To specifically assess the contribution of ERα, we next examined whether pharmacological blockade with MPP pharmacological inhibitor modified the stimulatory effect of sPIF on GSIS. In isolated mouse islets, sPIF increased insulin secretion under stimulatory glucose conditions, whereas this response was attenuated when sPIF was combined with MPP (Fig. 3C). Consistently, the fold change in insulin secretion was reduced in the presence of MPP (Fig. 3D), supporting the involvement of ERα in the functional response elicited by sPIF. These observations were further reproduced in MIN6 β-cells. After 24 h of treatment, sPIF enhanced GSIS, and this effect was attenuated by MPP (Fig. 3E). The same pattern was observed when insulin secretion was expressed as fold change (Fig. 3F). Together, these data indicate that sPIF enhances β-cell secretory responsiveness through a mechanism that requires ERα activity in both isolated mouse islets and MIN6 β-cells.

**Figure 3.**
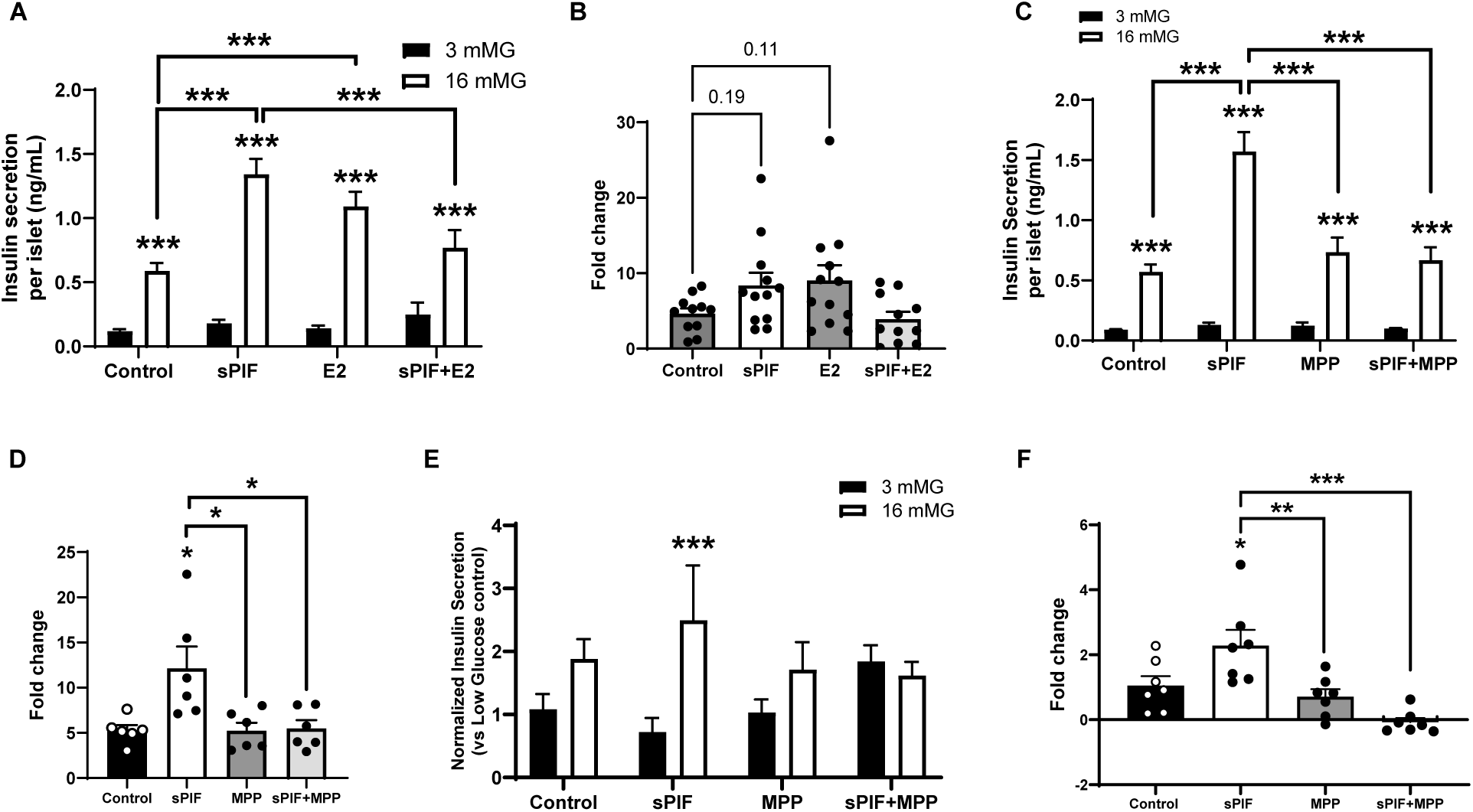
sPIF enhances GSIS through ERα-dependent signaling in mouse islets and MIN6 β-cells. **(A)** GSIS from isolated mouse islets treated for 24 h with vehicle, sPIF, E2 or the combination of sPIF and E2. **(B)** Fold change in insulin secretion calculated from the GSIS assay shown in panel A. 11-12 independent islet preparations from 3 independent experiments. **(C)** GSIS from isolated mouse islets treated for 24 h with sPIF in the absence or presence of MPP.**(D)** Fold change in insulin secretion calculated from the GSIS assay shown in panel C. 6 islet preparations from 2 independent experiments. **(E)** GSIS in MIN6 β-cells treated for 24 h with sPIF in the absence or presence of MPP. **(F)** Fold change in insulin secretion calculated from the MIN6 GSIS assay shown in panel E. 7 cell preparations from 2 independent experiments. Data are presented as mean ± SEM. Each dot represents one independent islet or cell preparation, as indicated. Statistical analysis was performed using two-way or one-way ANOVA followed by Bonferroni’s post hoc test. *p< 0.05, **p< 0.01, ***p<0.001

### 3.4 sPIF activates a kinase-mediated ERα signaling in MIN6 β-cells

To investigate the molecular basis of the adaptive effects induced by sPIF in β-cells, we first outlined a schematic model of the extranuclear-iniciated estrogen signaling pathway proposed to mediate its action (Fig. 4A). This model integrates membrane-initiated kinase activation with downstream convergence on ERα, providing a framework to analyze the signaling events triggered by sPIF. Western blot analysis in MIN6 β-cells after 24 h of treatment showed that sPIF increased phosphorylation of ERK, PKA substrates and CREB (Fig. 4B-C), indicating activation of signaling pathways associated with β-cell functional competence. In parallel, sPIF also enhanced phosphorylation of AKT and its downstream targets FOXO1 and GSK3β (Fig. 4D-E), consistent with activation of pathways involved in β-cell survival, growth and metabolic adaptation. Because these kinase pathways can converge on estrogen receptor signaling, we next analyzed ERα phosphorylation. sPIF increased ERα phosphorylation and elevated total ERα expression (Fig. 4F-G), supporting sustained engagement of estrogen receptor signaling within the kinase cascade activated by the peptide. Next, to determine whether these responses occurred rapidly, we analyzed the time-dependent activation of the signaling cascade after short-term sPIF exposure. Increased phosphorylation of key signaling mediators was already detectable within minutes, with activation observed at 5, 15 and 30 min after treatment (Fig. 4H-I). Together, these data indicate that sPIF activates a ERα-associated signaling pathway in β-cells, linking early kinase activation to ERα engagement and downstream pathways involved in β-cell function and proliferation.

**Figure 4.**
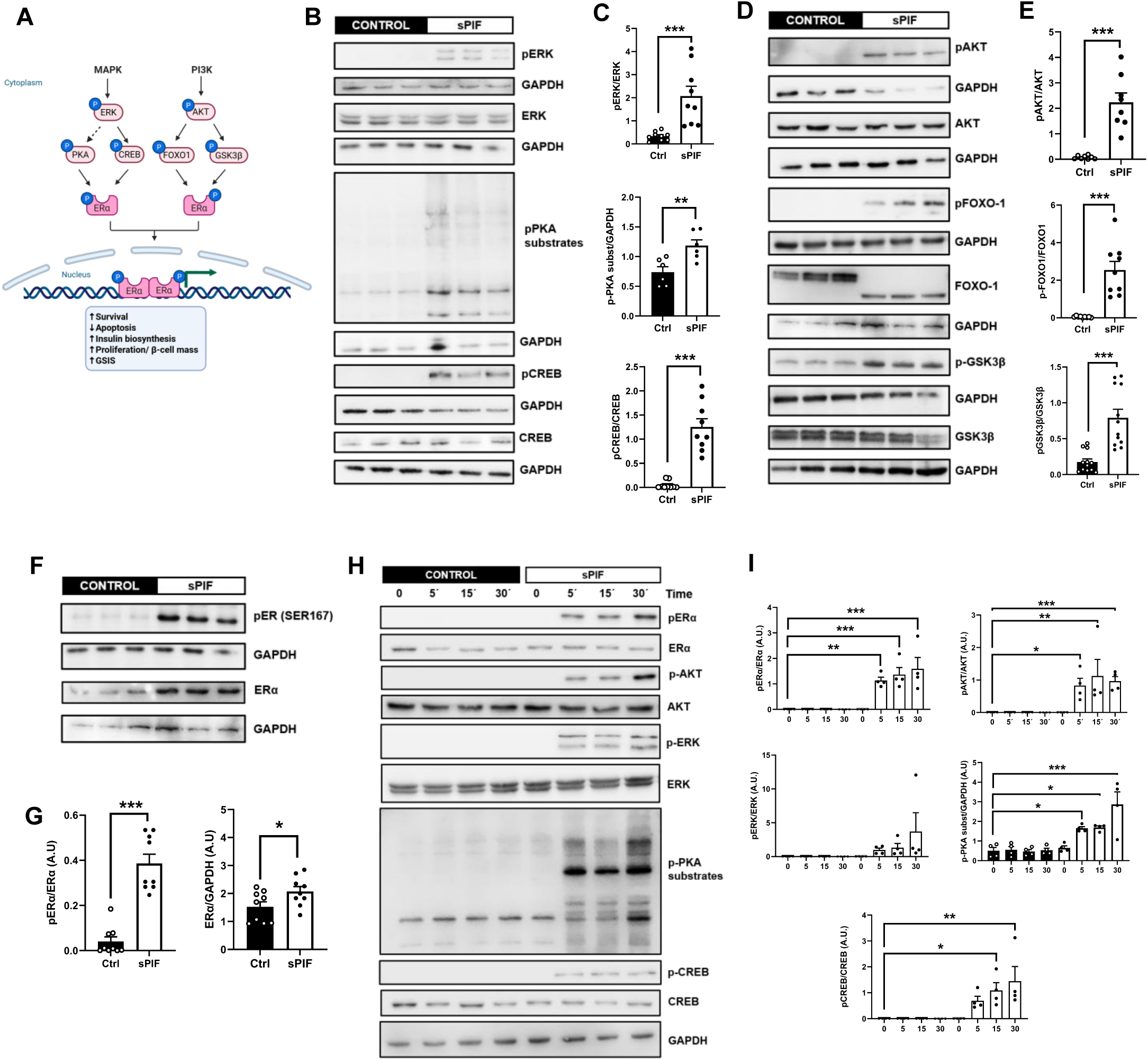
sPIF activates rapid non-canonical estrogen signaling in MIN6 β-cells. **(A)** Rapid non-canonical estrogen signal illustration **(B)** Representative Western blots showing phosphorylation of ERK, PKA substrates and CREB in MIN6 β-cells after 24 h of sPIF treatment.**(C)** Quantification of the Western blot data shown in panel B.**(D)** Representative Western blots showing phosphorylation of AKT and its downstream targets FOXO1 and GSK3β in MIN6 β-cells after 24 h of sPIF treatment.**(E)** Quantification of the Western blot data shown in panel D.**(F)** Representative Western blots showing phosphorylation of ERα and total ERα expression after 24 h of sPIF treatment.**(G)** Quantification of phospho-ERα and total ERα levels shown in panel F **(H)** Representative Western blots showing the time-dependent activation of the sPIF-induced signaling cascade at 5, 15 and 30 min after treatment.**(I)** Quantification of the time-course Western blot analysis shown in panel H. Data are presented as mean ± SEM. Each dot represents one well from three independent experiments for each experimental set. Unpaired t-test or one-Way ANOVA were applied. *p< 0.05, **p< 0.01, ***p<0.001.

### 3.5 ERα mediates and amplifies kinase-dependent signaling induced by sPIF

To define whether ERα contributes to the propagation of the rapid signaling response induced by sPIF, MIN6 β-cells were treated with sPIF in the absence or presence of the selective ERα antagonist MPP. Western blot analysis showed that MPP reduced the phosphorylation of key components of the sPIF-induced kinase cascade, including AKT, ERK and downstream signaling mediators (Fig. 5A-B). These data indicate that ERα activity contributes to the amplification of the signaling response triggered by sPIF. To further validate the requirement of ERα using a genetic approach, ERα expression was reduced in MIN6 β-cells by siRNA-mediated knockdown. Western blot analysis confirmed a significant decrease in ERα protein levels after silencing (Fig. 5C-D). Under these conditions, sPIF-induced phosphorylation of AKT and ERK was proportionally reduced in ERα-silenced cells compared with control siRNA-transfected cells (Fig. 5E-F). Together, these results support the idea that ERα is not only a downstream target of the sPIF-induced kinase cascade but also contributes to the propagation and stabilization of this signaling response in β-cells. This finding is consistent with a feed-forward mechanism in which ERα activity reinforces the rapid kinase signaling triggered by sPIF. Consistent with the effects observed with MPP, complementary pharmacological inhibition with ICI 182,780 similarly attenuated the sPIF-induced kinase response (Supplementary Fig. 2).

**Figure 5.**
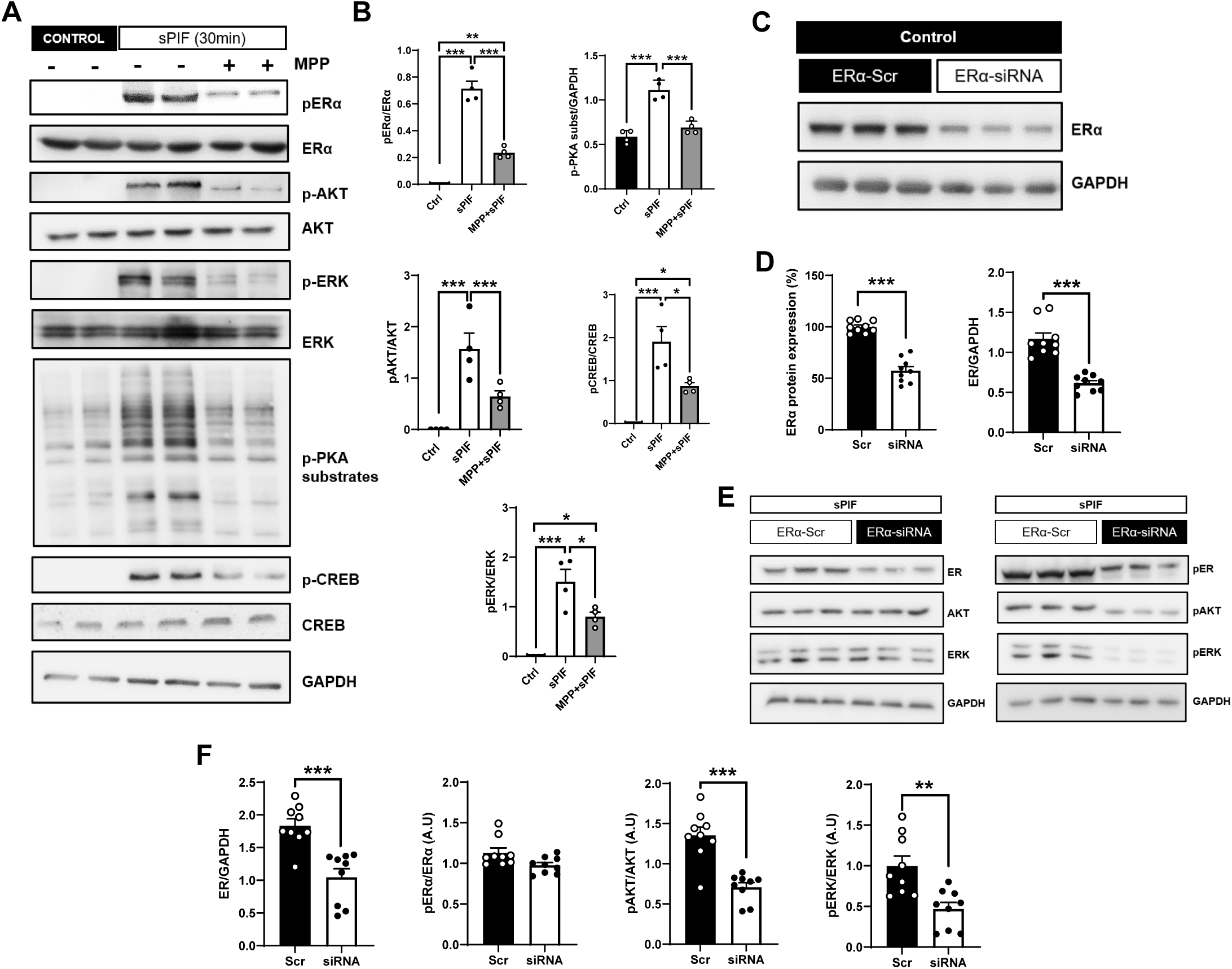
ERα contributes to the propagation of the rapid kinase response induced by sPIF. **(A)** Representative Western blots showing phosphorylation of the sPIF-induced kinase signaling cascade in MIN6 β-cells in the absence or presence of the selective ERα antagonist MPP.**(B)** Quantification of the Western blot data shown in panel A.**(C)** Representative Western blot showing ERα protein levels after siRNA-mediated ERα knockdown in MIN6 β-cells.**(D)** Quantification of ERα protein expression from the knockdown experiment shown in panel C.**(E)** Representative Western blots showing phosphorylation of AKT and ERK in control and ERα-silenced MIN6 β-cells treated with sPIF.**(F)** Quantification of the Western blot data shown in panel E. Data are presented as mean ± SEM. Each dot represents one well from 2-3 independent experiments performed in duplicate or triplicate. Statistical analysis was performed using unpaired Student’s t-test or one-way ANOVA followed by the appropriate post hoc test after assessment of normality, as indicated. *p< 0.05, **p< 0.01, ***p<0.001.

### 3.6 sPIF induces ERα phosphorylation and nuclear translocation through ERα-dependent signaling

To explore whether the kinase activation induced by sPIF was associated with changes in ERα localization and activation, we analyzed phospho-ERα and total ERα distribution by confocal microscopy in MIN6 β-cells and pancreatic sections from sPIF female treated-mice. Under control conditions, total ERα showed a predominantly cytoplasmic pattern, with low signal for phospho-ERα Ser118 and phospho-ERα Ser167. In contrast, sPIF treatment increased phospho-ERα signal at both residues and promoted nuclear accumulation of ERα (Fig. 6A). This response was reduced in the presence of MPP, indicating that ERα activity is required for the phosphorylation and nuclear redistribution induced by sPIF. Quantification of the confocal images confirmed an increase in phospho-ERα Ser118- and phospho-ERα Ser167-positive cells after sPIF treatment, together with increased fluorescence intensity for both phosphorylated forms (Fig. 6B). Orthogonal projections further supported the redistribution of ERα from the cytoplasmic compartment under basal conditions to the nuclear compartment after sPIF stimulation (Fig. 6C). Consistently, Pearson’s and Manders’ M1 coefficients showed increased nuclear localization of ERα in sPIF-treated cells (Fig. 6D). To determine whether this response was also present *in vivo*, pancreatic sections from mice treated with sPIF for 21 days were analyzed for phospho-ERα localization. In saline-treated mice, phospho-ERα Ser118 and Ser167 showed low nuclear colocalization within insulin-positive cells. In contrast, pancreatic islets from sPIF-treated mice displayed increased nuclear localization of phospho-ERα Ser118 (Fig. 6E-F) and phospho-ERα Ser167 (Fig. 6G-H). These data indicate that chronic sPIF exposure promotes sustained ERα phosphorylation and nuclear localization in pancreatic β-cells *in vivo*. Together, these findings support a model in which sPIF-induced kinase activation is coupled to phosphorylation-dependent nuclear translocation of ERα. This provides a mechanistic link between the rapid signaling events described above and the sustained functional and structural changes observed in β-cells after sPIF treatment.

**Figure 6.**
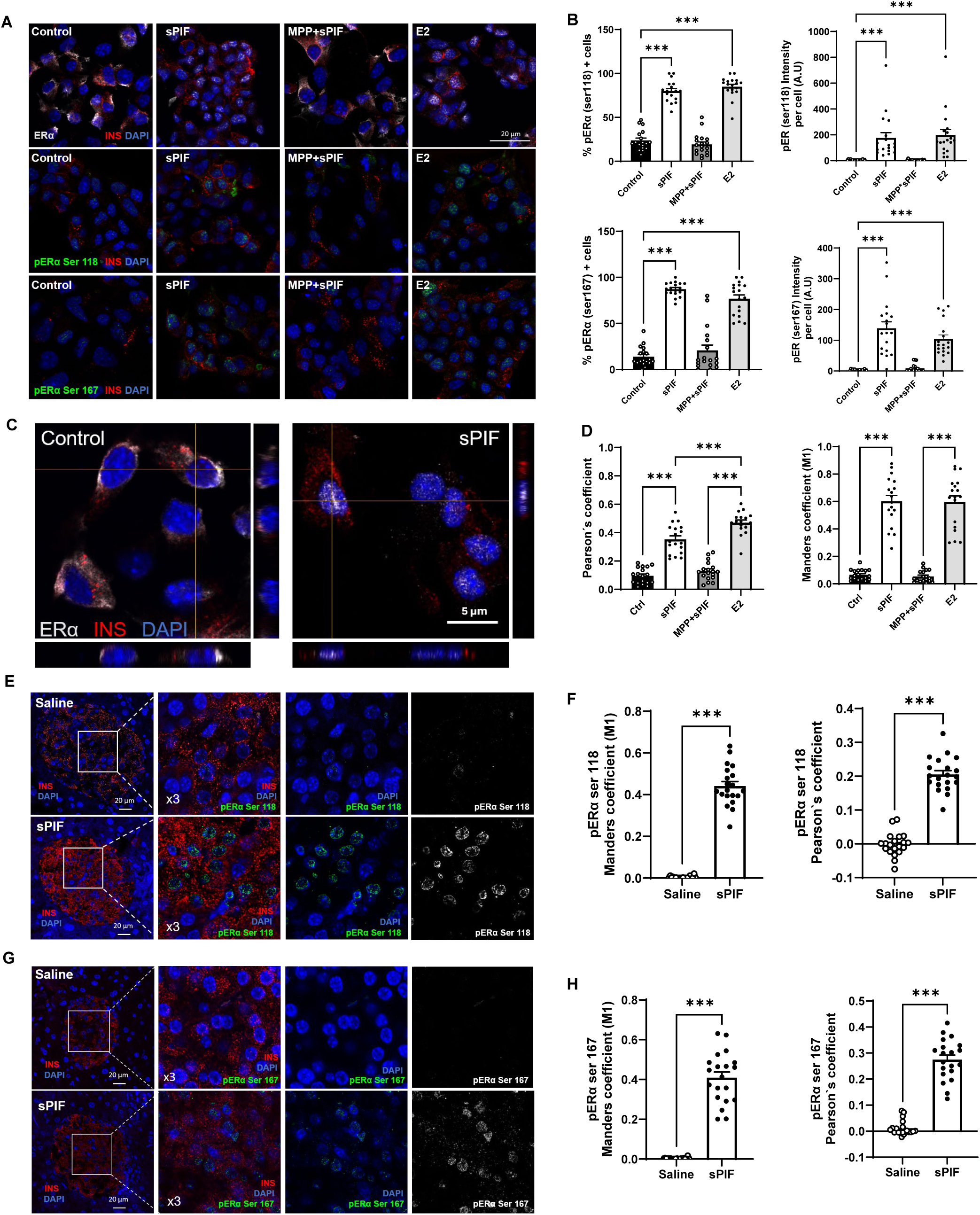
sPIF induces ERα-associated signaling and nuclear phospho-ERα translocation in β-cells. **(A)** Representative confocal immunofluorescence images of MIN6 β-cells stained for total ERα (white), phospho-ERα Ser118 (green), phospho-ERα Ser167(green), and insulin (red) under control conditions, sPIF treatment, MPP treatment and combined MPP+sPIF treatment.**(B)** Quantification of phospho-ERα Ser118- and phospho-ERα Ser167-positive cells and fluorescence intensity in the conditions shown in panel A.**(C)** Orthogonal projections showing Erα (white) subcellular localization in MIN6 β-cells under control and sPIF-treated conditions.**(D)** Quantification of ERα nuclear localization in MIN6 β-cells using Pearson’s correlation coefficient and Manders’ M1 coefficient.**(E)** Representative confocal immunofluorescence images of pancreatic sections from saline- and sPIF-treated mice after 21 days stained for phospho-ERα Ser118 (green) and insulin (red).**(F)** Quantification of phospho-ERα Ser118 nuclear localization in pancreatic β-cells using Pearson’s and Manders’ M1 coefficients.**(G)** Representative confocal immunofluorescence images of pancreatic sections from saline- and sPIF-treated mice after 21 days stained for phospho-ERα Ser167 (green) and insulin (red).**(H)** Quantification of phospho-ERα Ser167 nuclear localization in pancreatic β-cells using Pearson’s and Manders’ M1 coefficients. Data are presented as mean ± SEM. For MIN6 β-cells, each dot represents one randomly acquired field from two independent experiments performed in triplicate. For pancreatic sections, each dot represents one randomly acquired islet field from 3 mice pancreas per condition. Statistical analysis was performed using one-way ANOVA followed by the appropriate post hoc test or unpaired Student’s t-test after assessment of normality, as appropriate. *p< 0.05, **p< 0.01, ***p<0.001.

### 3.7 sPIF activates ERα-dependent signaling and enhances β-cell function in human islets

To assess the translational relevance of the signaling pathway identified, experiments were performed in isolated human islets from two women donors (Fig. 7A-B). Western blot analysis demonstrated that sPIF induced phosphorylation of ERK, AKT, CREB and PKA substrates, together with increased phosphorylation of ERα, reproducing the signaling cascade previously observed in MIN6 β-cells and mouse islets (Fig. 7A-B). Total protein levels remained unchanged, indicating that sPIF primarily modulates signaling through post-translational mechanisms. Functional assays showed that sPIF enhanced GSIS in woman islets, an effect that was significantly attenuated in the presence of the ERα antagonist MPP (Fig. 7C-D). Similarly, the fold change in insulin secretion was reduced by ERα inhibition. Together, these results indicate that the kinase-ERα signaling axis activated by sPIF is conserved in human β-cells and contributes to the regulation of insulin secretory responses.

**Figure 7.**
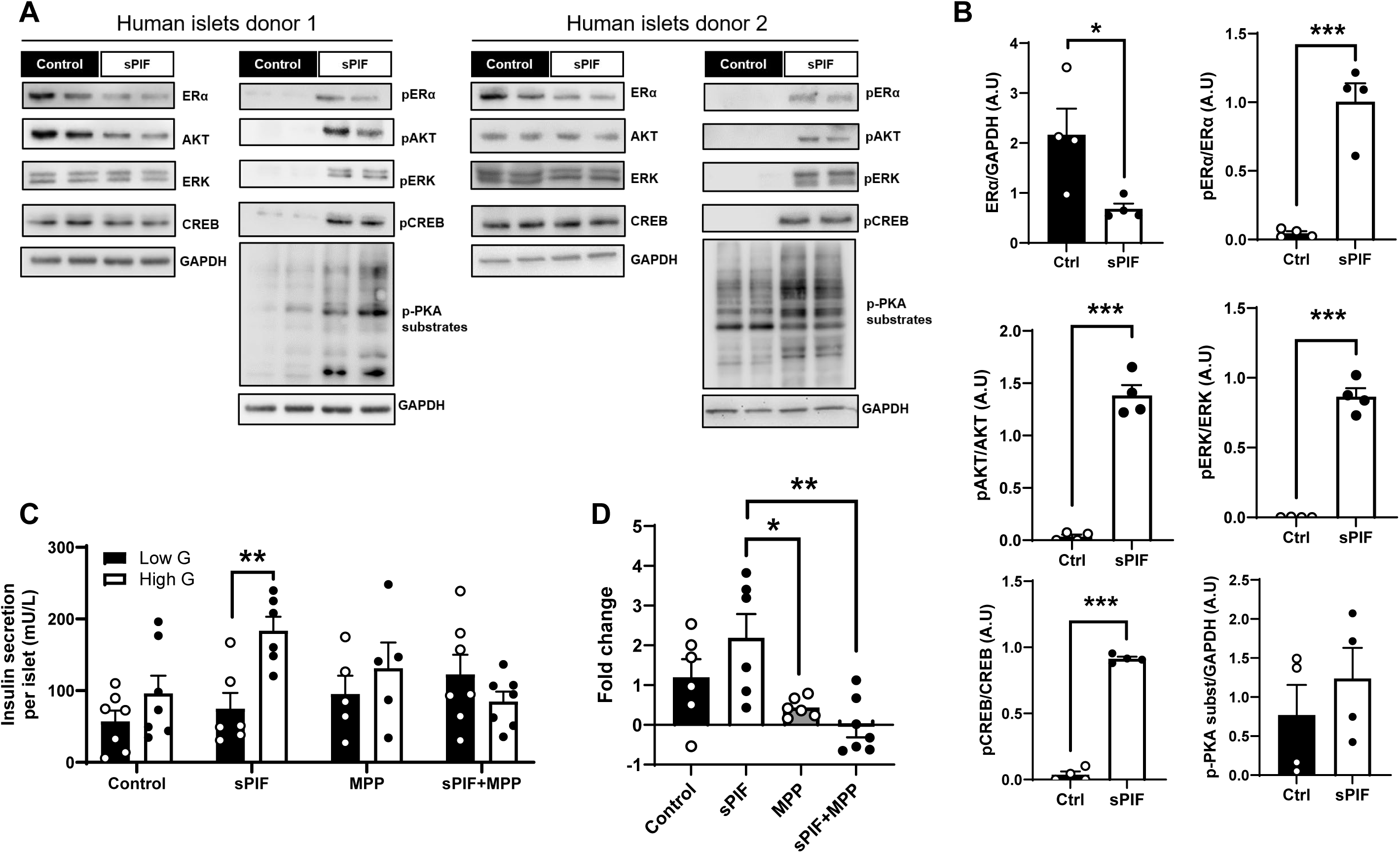
sPIF activates ERα-associated signaling and enhances GSIS in human islets. **(A)** Representative Western blots showing phosphorylation of ERα, AKT, ERK, CREB and PKA substrates in isolated human islets from two female donors under control conditions or after 24 h of sPIF treatment**.(B)** Quantification of the Western blot data shown in panel A.**(C)** GSIS from donor 2 human islets treated *ex vivo* with vehicle, sPIF, MPP or the combination of MPP+sPIF. **(D)** Fold change in insulin secretion calculated from the GSIS assay shown in panel C. Data are presented as mean ± SEM. For Western blot analyses, each dot represents one well from two independent experiments performed using islets from two female donors. For GSIS assays, each dot represents one independent islet preparation from donor 2. Statistical analysis was performed using unpaired Student’s t-test or one-way ANOVA followed by the appropriate post hoc test after assessment of normality, as indicated. *p< 0.05, **p< 0.01,***p<0.001.

## 4. Discussion

Our previous work demonstrated that sPIF enhances GSIS in MIN6 β-cells and human islets and improves glucose homeostasis and β-cell function in a murine model of diet-induced obesity [14]. In the present study, we extend these observations by placing PIF within the physiological context of maternal metabolic adaptation. Using a murine model of GDM, we found that pregnant females developed glucose intolerance and showed reduced circulating PIF levels at gestational days 8 and 15, with an overall decrease in PIF exposure across gestation. This finding is particularly relevant because day 15 coincides with the time point at which glucose intolerance was assessed, suggesting that plasmatic PIF levels are altered during a critical phase of maternal metabolic adaptation.

We next show that continuous sPIF exposure in non-pregnant female mice promotes a coordinated adaptive response in pancreatic β-cells that extends beyond a transient insulinotropic effect. This response followed a clear temporal sequence, with an early increase in β-cell proliferation after 1 week, enhanced circulating C-peptide levels in response to glucose after 15 days, and sustained improvement of *ex vivo* GSIS together with β-cell mass expansion after 21 days. This pattern is highly consistent with the physiological stages of gestational β-cell adaptation, in which early-to-mid gestation is characterized by increased β-cell proliferation, whereas β-cell mass expansion and sustained enhancement of insulin secretion become more evident at mid-to-late gestation [9–13, 24]. Moreover, the 21-day sPIF treatment window closely parallels the duration of murine gestation, further supporting that sPIF recapitulates key features of maternal β-cell adaptation to increased metabolic demand.

A key finding of this study is that islets isolated from female mice treated with sPIF for 21 days retained enhanced GSIS *ex vivo*. This indicates that sPIF does not act exclusively as an acute circulating insulin secretagogue but induces persistent functional changes within the islet. This observation is particularly relevant during pregnancy, when β-cell compensation requires both expansion of β-cell mass and a sustained increase in glucose-responsive insulin secretion [9, 13]. The enhanced *ex vivo* GSIS observed after chronic sPIF exposure therefore supports a role for this peptide in maintaining a compensated β-cell state. Moreover, the reduction in circulating PIF observed in GDM mice suggests that this response may be altered under gestational metabolic stress.

These findings extend the biological relevance previously attributed to sPIF beyond reproduction. In immune-mediated disease models, sPIF exerts systemic immunomodulatory effects without overt immune suppression and preserves pancreatic function in NOD mice, preventing diabetes development while maintaining islet architecture and insulin staining [25, 26]. Protective effects have also been reported in inflammatory and neurodegenerative experimental models [27, 28]. In addition, our previous work in diet-induced obesity established that sPIF improves β-cell function in a metabolic disease setting [14]. The present study extends this concept to the physiological increase in insulin demand associated with pregnancy by showing that sPIF promotes coordinated functional and structural changes in pancreatic β-cells. Thus, sPIF emerges not only as a protective and immunomodulatory peptide, but also as a regulator of β-cell plasticity under metabolic demand.

Because PIF is present in maternal circulation throughout gestation, its actions on β-cells occur within a complex hormonal environment rather than in isolation. E2 is a major gestational hormone whose circulating levels increase during pregnancy and whose signaling contributes to maternal metabolic and β-cell adaptation [13, 29]. We therefore examined sPIF in combination with E2 to model the physiological coexistence of these signals and to determine whether their insulinotropic actions were independent or convergent. Both sPIF and E2 enhanced GSIS after 24 h in female isolated islets, whereas their combined administration did not produce an additive response, suggesting convergence on shared regulatory mechanisms. This functional observation prompted us to investigate whether sPIF engages signaling pathways associated with estrogen action in β-cells.

To place our findings in a mechanistic context, previous studies have shown that E2 regulates complementary aspects of pancreatic β-cell biology through receptor-specific signaling pathways [23]. ERα increases insulin gene expression and intracellular insulin content through an estrogen response element-independent mechanism involving Src, ERK1/2 and NeuroD1 recruitment to the insulin promoter [17, 30]. ERβ, in turn, mediates the rapid insulinotropic response to E2 by promoting K_ATP_ channel closure and glucose-induced Ca²⁺ signaling [31], whereas GPER stimulates insulin secretion through EGFR transactivation, ERK activation and modulation by PI3K [32]. These receptors also contribute to β-cell protection against oxidative stress, inflammatory cytokines and lipotoxicity, and preserve islet lipid homeostasis under metabolic stress [33–35]. Together, these studies establish that estrogen signaling coordinates insulin biosynthesis, secretion, survival and metabolic homeostasis through distinct but interconnected mechanisms. This provides a relevant framework for our results, as sPIF activates ERK-, AKT- and PKA-dependent pathways and converges on ERα phosphorylation to promote β-cell function and adaptation. The ability of sPIF to engage kinase signaling is also supported by studies in other biological contexts. In decidual stromal cells, sPIF regulates the expression of implantation-related growth factor ligands through MAPK/ERK-dependent mechanisms, whereas in neuroprotective models it activates PKA/PKC signaling and promotes CREB phosphorylation [7,36]. Although these responses are context-dependent, they support the capacity of sPIF to modulate related kinase pathways across different cell types. Our findings extend this activity to the endocrine pancreas and identify ERα as the downstream integrative node linking sPIF-induced kinase activation to β-cell function.

Several lines of evidence from our study indicate that ERα is required for full propagation of the sPIF-induced signaling response. MPP attenuated the increase in GSIS induced by sPIF and reduced phosphorylation of AKT, ERK, PKA substrates, CREB and ERα. Moreover, siRNA-mediated knockdown of ERα led to a proportional decrease in sPIF-induced AKT and ERK activation. These data indicate that ERα is necessary for full signal propagation. ERα-contributes to β-cell survival and protection against oxidative, inflammatory and metabolic stress, and regulates signaling pathways including, ERK and PI3K/AKT activation, Ca²⁺ dynamics and insulin secretion [23,31–35]. To our knowledge, these findings provide the first evidence that sPIF induces ERα phosphorylation at Ser118 and Ser167 in pancreatic β-cells. This response was accompanied by nuclear accumulation of phosphorylated ERα both in MIN6 β-cells and in pancreatic β-cells from sPIF-treated mice, establishing a direct connection between the kinase cascade activated by sPIF and nuclear ERα signaling. Previous evidence for the phosphorylation-dependent regulation of these residues derives mainly from ERα-positive breast cancer models, where Ser118 and Ser167 integrate MAPK/ERK- and AKT-dependent inputs, respectively [37–39]. Our results therefore extend this mechanism to the pancreatic β-cell and identify ERα phosphorylation as a previously unrecognized component of the adaptive response induced by sPIF. Complementary data obtained with ICI 182,780 ER blocker are shown in the Supplemental Materials.

The phosphorylation of ERα at Ser118 and Ser167, together with its nuclear translocation, provides a mechanistic link between early cytosolic signaling and sustained transcriptional responses. These phosphorylation sites are known to integrate MAPK/ERK- and AKT-dependent inputs, respectively [37–39], and Ser118 phosphorylation has been related to promoter recruitment and gene-specific transcriptional activity of ERα [40]. Therefore, the coordinated phosphorylation and nuclear accumulation of ERα observed in our model support the idea that sPIF rapidly relays membrane-proximal signals to nuclear transcriptional responses involved in β-cell proliferation and functional adaptation. This interpretation is strengthened by the fact that nuclear phospho-ERα was observed both in MIN6 β-cells after sPIF treatment and in pancreatic β-cells from mice treated chronically with sPIF.

The relevance of this signaling axis is supported by our observations in isolated human islets. Previous studies have shown that estrogen signaling operates in human islets through non-canonical mechanisms involving extranuclear ERα and GPER, contributing to β-cell survival and rapid functional responses [33, 41]. In addition, rapid estrogenic regulation of insulin secretion has been reported in human islets, supporting the concept that human β-cells are responsive to membrane-associated estrogen receptor signaling [42]. In this context, our data add a new element by identifying sPIF, an embryo-derived peptide, as a pregnancy-associated input capable of activating a kinase-ERα signaling module in human islets. sPIF activated the same kinase pathways identified in murine β-cells and enhanced GSIS in human islets. Importantly, the stimulatory effect of sPIF on insulin secretion was lost in the presence of MPP, indicating that ERα activity is required for the functional response also in human β-cells. Thus, our findings do not simply extend the murine data to human islets but identify PIF as a previously unrecognized link between embryo-derived signaling and estrogen-associated β-cell regulation. Although the number of donors analyzed is limited, the conservation of both the biochemical response and the ERα-dependent secretory effect support the physiological relevance of this pathway in humans. This is particularly relevant in pregnancy, where human β-cells undergo functional adaptation and where inadequate β-cell compensation contributes to GDM [9, 10].

Taken together, our findings identify PIF as a pregnancy-associated signal linking embryo-derived cues to maternal β-cell adaptation. Sustained sPIF exposure recapitulated the temporal progression of gestational β-cell compensation, from early proliferation to increased secretory capacity and expansion of β-cell mass. The reduction in circulating PIF observed in the GDM model further indicates that this endogenous signal is altered under maternal metabolic stress. These results establish a mechanistic and physiological link between PIF availability and the capacity of maternal β-cells to meet gestational insulin demand, suggesting that disruption of this adaptive axis may contribute to insufficient β-cell compensation in GDM.

These findings position PIF as a previously unrecognized embryo-to-maternal-pancreas signal, distinct from classical placental and maternal hormonal cues, that may act as a missing link between early embryonic signaling and maternal β-cell compensation during pregnancy.

## Acknowledgements

The authors gratefully acknowledge Dr. Ángel Nadal and Dr. Iván Quesada from Universidad Miguel Hernández, and Dr. Sergi Soriano and Universidad de Alicante for their scientific assistance and constructive discussions. The authors also thank Dr. Diego Sánchez and Dr. Maria Dolores Ganfornina from University of Valladolid for their support throughout this work. Thanks also to Borja Santirso for his technical work.

## Funding

This work was supported by grants PID2023-150827OA-I00 to BM; PID2022-136605OB-C21 to IC; PID2022-136605OB-C22 to GP; and PID2023-146795OB-I00 to PA-M, funded by MICIU/AEI/10.13039/501100011033 and, where applicable, by “ERDF A way of making Europe”. M.L.C was supported by a CIBER BBN contract funded by Instituto de Salud Carlos III (ISCIII). This work was also supported by grant VA119P24 to IC and BM, funded by the Junta de Castilla y León.

### Author contributions

R.P.-M. performed most of the experimental work, contributed to conceptualization, data interpretation and scientific discussion. A.S.-Y. contributed to experimental work, data analysis and manuscript revision. S.H.-d.l.R. contributed to experimental development, data acquisition and manuscript revision. T.B.-B. contributed to experimental work. M.A.-M., M.M.-M. and M.D.M provided technical assistance. M.R. and M.L.C performed sPIF synthesis. N.P. and P.M. developed and provided the anti-sPIF antibody. C.B. and J.S. provided and isolated human islets. G.P. contributed to conceptualization and manuscript revision. I.C.-C. contributed to experimental development, conceptualization, data interpretation and manuscript revision. P.A.-M. contributed to experimental development, technical work, conceptualization, scientific discussion and manuscript revision. B.M. conceived the study, formulated the hypothesis, supervised and organized the project, contributed to experimental work, data interpretation and conceptualization, and wrote the manuscript. All authors reviewed and approved the final version of the manuscript.

### Declaration of generative AI and AI-assisted technologies

AI-assisted tools were used only to support language editing, manuscript organization and refinement of conceptual descriptions. These tools were not used to generate the scientific content of the manuscript, design the study, analyze data, interpret results or draw conclusions. All scientific content, data interpretation and conclusions were developed, reviewed and approved by the authors. No AI tools were used to generate, modify or enhance figures, graphical abstract, Western blots, immunofluorescence, histology, confocal microscopy images or any other primary experimental data.

## Data availability

Data is available from the corresponding authors upon reasonable request.

## References

1. Roussev RG, Barnea ER, Thomason EJ, Coulam CB. A novel bioassay for detection of preimplantation factor (PIF). Am J Reprod Immunol. 1995;33:68–73. PMID: 7619236.

2. Coulam CB, Roussev RG, Thomason EJ, Barnea ER. Preimplantation factor (PIF) predicts subsequent pregnancy loss. Am J Reprod Immunol. 1995;34:88–92. PMID: 8526994.

3. Roussev RG, Coulam CB, Barnea ER. Development and validation of an assay for measuring preimplantation factor (PIF) of embryonal origin. Am J Reprod Immunol. 1996;35:281–287. PMID: 8962662

4. Barnea ER, Simon J, Levine SP, Coulam CB, Taliadouros GS, Leavis PC. Progress in characterization of pre-implantation factor in embryo cultures and in vivo. Am J Reprod Immunol. 1999;42:95–99. PMID: 10476691.

5. Stamatkin CW, Roussev RG, Stout M, Absalon-Medina V, Ramu S, Goodman C, et al. PreImplantation Factor (PIF) correlates with early mammalian embryo development-bovine and murine models. Reprod Biol Endocrinol. 2011;9:63. PMID: 21569635

6. Paidas MJ, Krikun G, Huang SJ, Jones R, Romano M, Annunziato J, et al. A genomic and proteomic investigation of the impact of preimplantation factor on human decidual cells. Am J Obstet Gynecol. 2010;202:459.e1–459.e8. PMID: 20452489

7. Barnea ER, Kirk D, Paidas MJ. PreImplantation Factor (PIF) promoting role in embryo implantation: increases endometrial Integrin-α2β3, amphiregulin and epiregulin while reducing betacellulin expression via MAPK in decidua. Reprod Biol Endocrinol. 2012;10:50. PMID: 22788113

8. O’Brien CB, Barnea ER, Martin P, Levy C, Sharabi E, Bhamidimarri KR, et al. Randomized, double-blind, placebo-controlled, single ascending dose trial of synthetic preimplantation factor in autoimmune hepatitis. Hepatol Commun. 2018;2:1235–1246. PMID: 30411073.

9. Pretorius M, Huang C. Beta-Cell Adaptation to Pregnancy – Role of Calcium Dynamics. Front Endocrinol (Lausanne). 2022;13:853876. PMID: 35399944.

10. Butler AE, Cao-Minh L, Galasso R, Rizza RA, Corradin A, Cobelli C, Butler PC. Adaptive changes in pancreatic beta cell fractional area and beta cell turnover in human pregnancy. Diabetologia. 2010 Oct;53(10):2167–76. doi: 10.1007/s00125-010-1809-6. Epub 2010 Jun 5. PMID: 20523966; PMCID: PMC2931643.

11. Seedat F, Holden K, Davis S, Fischer R, Bancroft J, Drydale E, Kandzija N, Todd JA, Vatish M, Stefana MI. A new paradigm of islet adaptations in human pregnancy: insights from immunohistochemistry and proteomics. Nat Commun. 2025 Jul 21;16(1):6687. doi: 10.1038/s41467-025-61852-5. PMID: 40691142; PMCID: PMC12280027.

12. Usman TO, Chhetri G, Yeh H, Dong HH. Beta-cell compensation and gestational diabetes. J Biol Chem. 2023;299:105405. PMID: 38229396.

13. Moyce BL, Dolinsky VW. Maternal β-cell adaptations in pregnancy and placental signalling: implications for gestational diabetes. Int J Mol Sci. 2018;19:3467. PMID: 30400566.

14. Sanz-González A, Cózar-Castellano I, Broca C, Sabatier J, Acosta GA, Royo M, et al. Pharmacological activation of insulin-degrading enzyme improves insulin secretion and glucose tolerance in diet-induced obese mice. Diabetes Obes Metab. 2023;25:3268–3278. PMID: 37493025

15. Boronat-Belda T, Ferrero H, Soriano S, Ribes-García E, Betoret-Gustems R, Martínez-Bañón D, et al. Increased TGF-β/Activin-Smad2 signaling is associated with pancreatic β-cell dysfunction and glucose intolerance in gestational diabetes mellitus. Mol Metab. 2026;103:102274. PMID: 41115652.

16. Alonso-Magdalena P, Morimoto S, Ripoll C, Fuentes E, Nadal A. The estrogenic effect of bisphenol A disrupts pancreatic beta-cell function in vivo and induces insulin resistance. Environ Health Perspect. 2006;114:106–112. PMID: 16393666.

17. Alonso-Magdalena P, Ropero AB, Carrera MP, Cederroth CR, Baquié M, Gauthier BR, Nef S, Stefani E, Nadal A. Pancreatic insulin content regulation by the estrogen receptor ER alpha. PLoS One. 2008 Apr 30;3(4):e2069. doi: 10.1371/journal.pone.0002069. PMID: 18446233; PMCID: PMC2323613.

18. Villar-Pazos S, Martinez-Pinna J, Castellano-Muñoz M, Alonso-Magdalena P, Marroqui L, Quesada I, Gustafsson JA, Nadal A. Molecular mechanisms involved in the non-monotonic effect of bisphenol-a on ca2+ entry in mouse pancreatic β-cells. Sci Rep. 2017 Sep 18;7(1):11770. doi: 10.1038/s41598-017-11995-3. Erratum in: Sci Rep. 2018 Mar 6;8(1):4262. doi: 10.1038/s41598-018-21309-w. PMID: 28924161; PMCID: PMC5603522.

19. Babiloni-Chust I, Dos Santos RS, Medina-Gali RM, Perez-Serna AA, Encinar JA, Martinez-Pinna J, Gustafsson JA, Marroqui L, Nadal A. G protein-coupled estrogen receptor activation by bisphenol-A disrupts the protection from apoptosis conferred by the estrogen receptors ERα and ERβ in pancreatic beta cells. Environ Int. 2022 Jun;164:107250. doi: 10.1016/j.envint.2022.107250. Epub 2022 Apr 19. PMID: 35461094.

20. Merino B, Casanueva-Álvarez E, Quesada I, González-Casimiro CM, Fernández-Díaz CM, Postigo-Casado T, Leissring MA, Kaestner KH, Perdomo G, Cózar-Castellano I. Insulin-degrading enzyme ablation in mouse pancreatic alpha cells triggers cell proliferation, hyperplasia and glucagon secretion dysregulation. Diabetologia. 2022 Aug;65(8):1375–1389. doi: 10.1007/s00125-022-05729-y. Epub 2022 Jun 2. Erratum in: Diabetologia. 2024 Jan;67(1):218-219. doi: 10.1007/s00125-023-06037-9. PMID: 35652923; PMCID: PMC9283140.

21. Ballesteros B, Barceló D, Camps F, Marco MP. Preparation of antisera and development of a direct enzyme-linked immunosorbent assay for the determination of the antifouling agent Irgarol 1051. Anal Chim Acta. 1997;347:139–147. DOI: 10.1016/S0003-2670(97)00317-6.

22. Merino B, Fernández-Díaz CM, Parrado-Fernández C, González-Casimiro CM, Postigo-Casado T, Lobatón CD, et al. Hepatic insulin-degrading enzyme regulates glucose and insulin homeostasis in diet-induced obese mice. Metabolism. 2020;113:154352. PMID: 32916153

23. Nadal A, Alonso-Magdalena P, Soriano S, Ripoll C, Fuentes E, Quesada I, et al. Role of estrogen receptors alpha, beta and GPER1/GPR30 in pancreatic beta-cells. Front Biosci (Landmark Ed). 2011;16:251–260. PMID: 21196169

24. Ikushima YM, Awazawa M, Kobayashi N, Osonoi S, Takemiya S, Kobayashi N, et al. MEK/ERK signaling in β-cells bifunctionally regulates β-cell mass and glucose-stimulated insulin secretion response to maintain glucose homeostasis. Diabetes. 2021;70:1519–1535. PMID: 33906910

25. Weiss L, Bernstein S, Jones R, Amunugama R, Krizman D, Jebailey L, Almogi-Hazan O, Yekhtin Z, Shiner R, Reibstein I, Triche E, Slavin S, Or R, Barnea ER. Preimplantation factor (PIF) analog prevents type I diabetes mellitus (TIDM) development by preserving pancreatic function in NOD mice. Endocrine. 2011 Aug;40(1):41–54. doi: 10.1007/s12020-011-9438-5. Epub 2011 Mar 22. Erratum in: Endocrine. 2011 Aug;40(1):55. Hazan, Osnat [corrected to Almogi-Hazan, Osnat]; Yachtin, Janna [corrected to Yekhtin, Zhanna]. PMID: 21424847.

26. Barnea ER. Applying embryo-derived immune tolerance to the treatment of immune disorders. Ann N Y Acad Sci. 2007 Sep;1110:602–18. doi: 10.1196/annals.1423.064. PMID: 17911476.

27. Barnea ER, Kirk D, Ramu S, Rivnay B, Roussev R, Paidas MJ. PreImplantation Factor (PIF) orchestrates systemic antiinflammatory response by immune cells: effect on peripheral blood mononuclear cells. Am J Obstet Gynecol. 2012 Oct;207(4):313.e1-11. doi: 10.1016/j.ajog.2012.07.017. PMID: 23021695.

28. Hayrabedyan S, Shainer R, Yekhtin Z, Weiss L, Almogi-Hazan O, Or R, et al. Synthetic PreImplantation Factor (sPIF) induces posttranslational protein modification and reverses paralysis in EAE mice. Sci Rep. 2019;9:12869. PMID: 31578341.

29. Parisi F, Fenizia C, Introini A, Zavatta A, Scaccabarozzi C, Biasin M, Savasi V. The pathophysiological role of estrogens in the initial stages of pregnancy: molecular mechanisms and clinical implications for pregnancy outcome from the periconceptional period to end of the first trimester. Hum Reprod Update. 2023 Nov 2;29(6):699–720. doi: 10.1093/humupd/dmad016. PMID: 37353909; PMCID: PMC10628507.

30. Wong WP, Tiano JP, Liu S, Hewitt SC, Le May C, Dalle S, Katzenellenbogen JA, Katzenellenbogen BS, Korach KS, Mauvais-Jarvis F. Extranuclear estrogen receptor-alpha stimulates NeuroD1 binding to the insulin promoter and favors insulin synthesis. Proc Natl Acad Sci U S A. 2010 Jul 20;107(29):13057–62. doi: 10.1073/pnas.0914501107. Epub 2010 Jun 29. PMID: 20616010; PMCID: PMC2919966.

31. Soriano S, Ropero AB, Alonso-Magdalena P, Ripoll C, Quesada I, Gassner B, Kuhn M, Gustafsson JA, Nadal A. Rapid regulation of K(ATP) channel activity by 17{beta}-estradiol in pancreatic {beta}-cells involves the estrogen receptor {beta} and the atrial natriuretic peptide receptor. Mol Endocrinol. 2009 Dec;23(12):1973–82. doi: 10.1210/me.2009-0287. Epub 2009 Oct 23. PMID: 19855088; PMCID: PMC5419133.

32. Sharma G, Prossnitz ER. Mechanisms of estradiol-induced insulin secretion by the G protein-coupled estrogen receptor GPR30/GPER in pancreatic β-cells. Endocrinology. 2011;152:3030–3039. PMID: 21673097.

33. Le May C, Chu K, Hu M, Ortega CS, Simpson ER, Korach KS, Tsai MJ, Mauvais-Jarvis F. Estrogens protect pancreatic beta-cells from apoptosis and prevent insulin-deficient diabetes mellitus in mice. Proc Natl Acad Sci U S A. 2006 Jun 13;103(24):9232–7. doi: 10.1073/pnas.0602956103. Epub 2006 Jun 5. PMID: 16754860; PMCID: PMC1482595.

34. Liu S, Le May C, Wong WPS, Ward RD, Clegg DJ, Marcelli M, et al. Importance of extranuclear estrogen receptor-alpha and membrane G protein-coupled estrogen receptor in pancreatic islet survival. Diabetes. 2009;58:2292–2302. PMID: 19587358.

35. Tiano JP, Delghingaro-Augusto V, Le May C, Liu S, Kaw MK, Khuder SS, Latour MG, Bhatt SA, Korach KS, Najjar SM, Prentki M, Mauvais-Jarvis F. Estrogen receptor activation reduces lipid synthesis in pancreatic islets and prevents β cell failure in rodent models of type 2 diabetes. J Clin Invest. 2011 Aug;121(8):3331–42. doi: 10.1172/JCI44564. Epub 2011 Jul 11. PMID: 21747171; PMCID: PMC3148728.

36. Mueller M, Schoeberlein A, Zhou J, Joerger-Messerli M, Oppliger B, Reinhart U, et al. PreImplantation Factor bolsters neuroprotection via modulating Protein Kinase A and Protein Kinase C signaling. Cell Death Differ. 2015;22:2078–2086. PMID: 25976303.

37. Chen D, Washbrook E, Sarwar N, Bates GJ, Pace PE, Thirunuvakkarasu V, et al. Phosphorylation of human estrogen receptor alpha at serine 118 by two distinct signal transduction pathways revealed by phosphorylation-specific antisera. Oncogene. 2002;21:4921–4931. PMID: 12118371.

38. Jiang J, Sarwar N, Peston D, Kulinskaya E, Shousha S, Coombes RC, et al. Phosphorylation of estrogen receptor-alpha at Ser167 is indicative of longer disease-free and overall survival in breast cancer patients. Clin Cancer Res. 2007;13:5769–5776. PMID: 17908967.

39. Yamashita H, Nishio M, Toyama T, Sugiura H, Kondo N, Kobayashi N, et al. Low phosphorylation of estrogen receptor alpha (ERalpha) serine 118 and high phosphorylation of ERalpha serine 167 improve survival in ER-positive breast cancer. Endocr Relat Cancer. 2008;15:755–763. PMID: 18550720.

40. Duplessis TT, Williams CC, Hill SM, Rowan BG. Phosphorylation of Estrogen Receptor α at serine 118 directs recruitment of promoter complexes and gene-specific transcription. Endocrinology. 2011;152:2517–2526. PMID: 21505052.

41. Liu S, Mauvais-Jarvis F. Rapid, nongenomic estrogen actions protect pancreatic islet survival. Islets. 2009 Nov-Dec;1(3):273–5. doi: 10.4161/isl.1.3.9781. PMID: 20634925; PMCID: PMC2903892.

42. Soriano S, Alonso-Magdalena P, García-Arévalo M, Novials A, Muhammed SJ, Salehi A, Gustafsson JA, Quesada I, Nadal A. Rapid insulinotropic action of low doses of bisphenol-A on mouse and human islets of Langerhans: role of estrogen receptor β. PLoS One. 2012;7(2):e31109. doi: 10.1371/journal.pone.0031109. Epub 2012 Feb 8. PMID: 22347437; PMCID: PMC3275611.

