## Supplementary material for "Preimplantation factor (PIF) links embryo-derived signaling to maternal pancreatic β-cell adaptation through an ERα-dependent pathway": Supplemenry tables and figures.pdf

### Supplemental figures

Pascua-Maestro et al. 2026

#### Supplementary. Table 1

| Target protein | Antibody Supplier | Catalog number | Host specie | Working dilution | Experimental conditions | Method |
|---|---|---|---|---|---|---|
| **PIF** | ICTS NANBIOSIS – IQAC | See methods | Rabbit | 1:2000 | ON, 4ºC | DB |
| **Rb** | Cell Signaling | 9309S | Rabbit | 1:1000 | ON, 4ºC | WB |
| **pRb S807** | Cell Signaling | 9308S | Rabbit | 1:1000 | ON, 4ºC | WB |
| **pERK 1/2 (p44/42 MAPK)** | Cell Signaling | 9101 | Rabbit | 1:1000 | ON, 4ºC | WB |
| **ERK 1/2 (p44/42 MAPK)** | Cell Signaling | 9102 | Rabbit | 1:1000 | ON, 4ºC | WB |
| **pCREB** | Cell Signaling | 9198S | Rabbit | 1:2000 | ON, 4ºC | WB |
| **CREB** | Cell Signaling | 9197S | Rabbit | 1:2000 | ON, 4ºC | WB |
| **pAKT** | Cell Signaling | 13038 | Rabbit | 1:1000 | ON, 4ºC | WB |
| **AKT** | Cell Signaling | 9272S | Rabbit | 1:1000 | ON, 4ºC | WB |
| **FOXO-1** | Cell Signaling | 2880S | Rabbit | 1:1000 | ON, 4ºC | WB |
| **pFOXO-1** | Cell Signaling | 9461S | Rabbit | 1:1000 | ON, 4ºC | WB |
| **pGSK3β** | Cell Signaling | 9327S | Rabbit | 1:1000 | ON, 4ºC | WB |
| **GSK3β** | Cell Signaling | 5676S | Rabbit | 1:1000 | ON, 4ºC | WB |
| **pERα** | Invitrogen | PA5-99347 | Rabbit | 1:1000 | ON, 4ºC | WB |
| **ERα** | Cell Signaling | 13258 | Rabbit | 1:1000 | ON, 4ºC | WB |
| **pPKA substrates** | Cell Signaling | 9624 | Rabbit | 1:50000 | ON, 4ºC | WB |
| **GAPDH** | Sigma Aldrich | G9545 | Mouse | 1:40000 | 1 h, RT | WB |
| **Anti-mouse IgG H&L (HRP)** | Abcam | Ab6789 | Goat | 1: 5000 | 30 min, RT | WB |
| **Anti-Rabbit** | Jackson Immunoresearch | 711-035-152 | Goat | 1:20000 | 30 min, RT | WB/DB |
| **p84** | Abcam | ab487 | Mouse | 1:5000 | 1 hour, RT | IF |
| **Glucagon** | Abcam | ab30788 | Rabbit | 1:2000 | ON, 4ºC | IF |
| **Somatostatin** | Abcam | ab30788 | Rat | 1:250 | ON, 4ºC | IF |
| **Insulin** | Sigma-Aldrich | I2018 | Mouse | 1:1000 | ON, 4ºC | IF |
| **Ki67** | Invitrogen | MA5-14520 | Rabbit | 1:500 | ON, 4ºC | IF |
| **pERα Ser 118** | Invitrogen | PA5-99347 | Rabbit | 1:200 | ON, 4ºC | IF |
| **pERα Ser 167** | Invitrogen | PA5-37570 | Rabbit | 1:200 | ON, 4ºC | IF |
| **ERα** | Invitrogen | MA1-80216 | Mouse | 1:200 | ON, 4ºC | IF |
| **Alexa Fluor 488** | Invitrogen | A11001 | Goat | 1:1000 | 30 min, RT | IF |
| **Alexa Fluor 594** | Invitrogen | A11005 | Goat | 1:1000 | 30 min, RT | IF |

**Supplementary Table 1. Antibodies used for Dot Blot, Western blot and immunofluorescence analyses.** The table lists the target protein, antibody supplier, catalogue number, host specie, working dilution and experimental conditions used for each primary and secondary antibody. DB, dot blot; WB, Western blot; IF, immunofluorescence; RT, room temperature; ON, overnight.

#### Supplementary. Fig 1

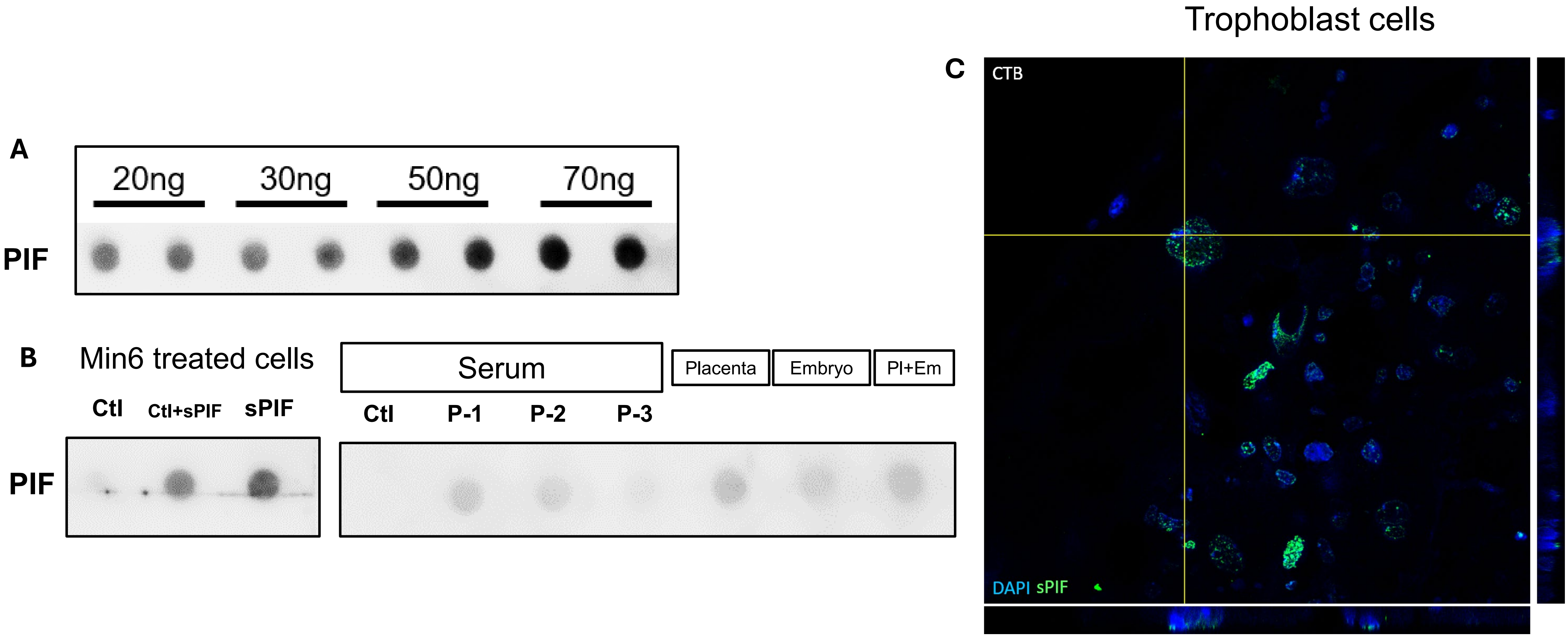


**Supplementary Figure 1. Validation of the anti-PIF antibody for detection of synthetic and endogenous PIF.(A)** Representative dot blot showing antibody detection of increasing concentrations of synthetic PIF.**(B)** Dot blot analysis of PIF detection in different biological and experimental samples, including untreated MIN6 cell lysate, MIN6 cell lysate supplemented with synthetic PIF, synthetic PIF alone, serum from non-pregnant control mice, sera from pregnant mice, and samples obtained from placenta, embryo and fetoplacental unit.**(C)** Representative immunofluorescence images of placental tissue showing PIF-positive signal in trophoblast cells (green).Dot blot and immunofluorescence analyses were performed to validate the specificity and sensitivity of the antibody for the detection of both synthetic PIF and endogenous PIF-containing samples. Images are representative of the antibody validation experiments.

Supplementary. Fig 2

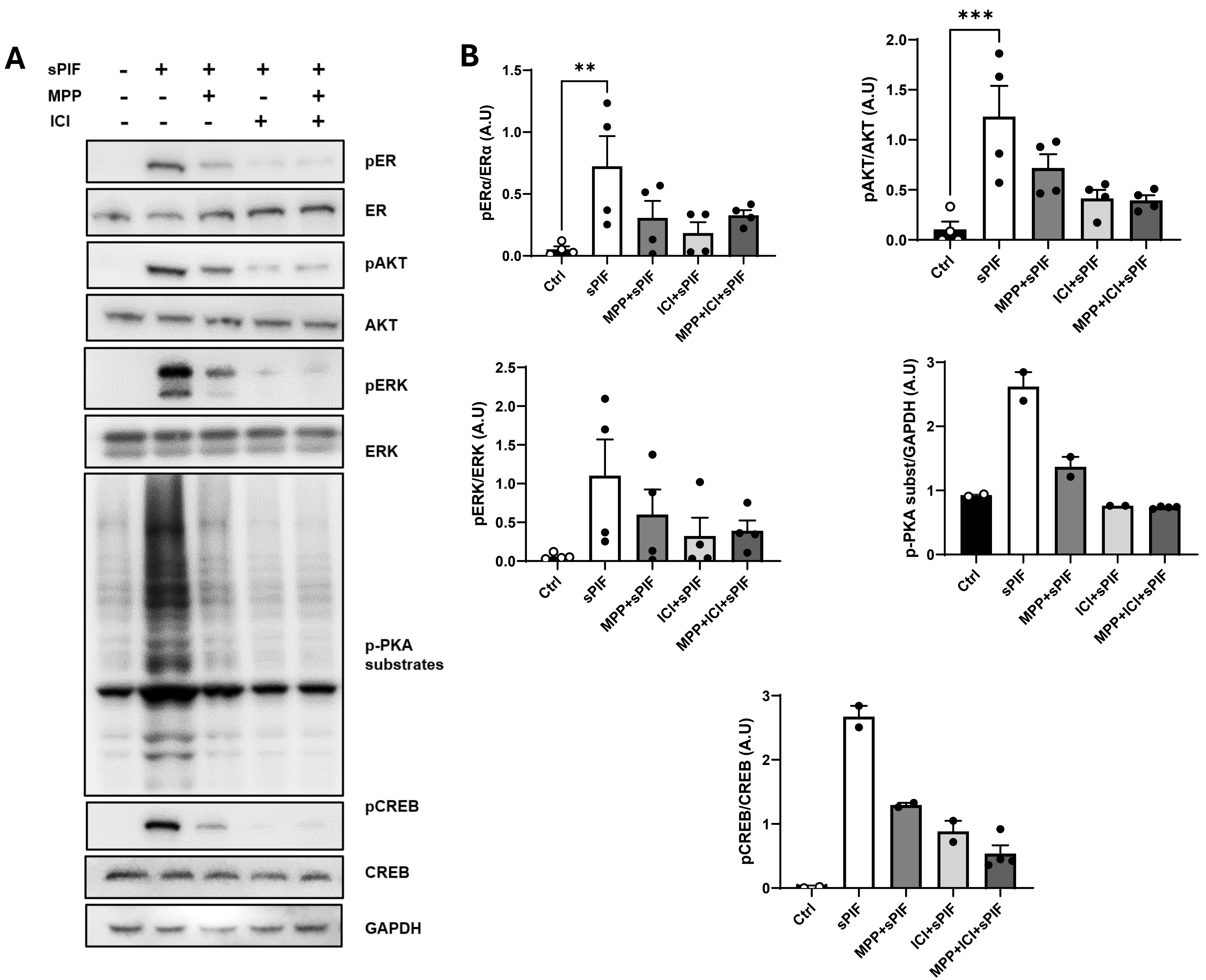

**Supplementary Figure 2**. Estrogen receptor inhibition attenuates the sPIF-induced kinase signaling cascade in MIN6 β-cells.**(A)** Representative Western blots showing the phosphorylation of ERα, AKT, ERK, PKA substrates and CREB in MIN6 β-cells treated with sPIF in the absence or presence of MPP, ICI 182,780, or the combination of both inhibitors.**(B)** Quantification of phospho-ERα, phospho-AKT, phospho-ERK, phospho-PKA substrates and phospho-CREB levels from the Western blots shown in panel A. Pharmacological inhibition of estrogen receptor signaling reduced the sPIF-induced kinase response, with a pattern comparable to that observed with MPP in the main experimental series. Data are presented as mean ± SEM. Each dot represents one well from two independent experiments performed in single or duplicate wells, as indicated. Statistical analysis was performed using one-way ANOVA followed by the appropriate post hoc test. *$p < 0.05$, **$p < 0.01$, ***$p < 0.001$.
